# NT-CRISPR: Combining natural transformation and CRISPR/Cas9 counterselection for markerless and scarless genome editing in *Vibrio natriegens*

**DOI:** 10.1101/2021.08.02.454823

**Authors:** Daniel Stukenberg, Josef Hoff, Anna Faber, Anke Becker

**Affiliations:** Center for Synthetic Microbiology, Philipps-Universität Marburg, Marburg, Germany; Max-Planck Institute for Terrestrial Microbiology, Marburg, Germany

## Abstract

The fast-growing bacterium *Vibrio natriegens* has recently gained increasing attention as a novel chassis organism for a wide range of projects. To fully harness the potential of this fascinating bacterium, convenient and highly efficient genome editing methods are indispensable to create novel strains, tailored for specific applications. *V. natriegens* is able to take up free DNA and incorporate it into its genome by homologous recombination. This process, called natural transformation, was tamed for genome editing. It displays a high efficiency and is able to mediate uptake of multiple DNA fragments, thereby allowing multiple simultaneous edits. Here, we describe NT-CRISPR, a combination of natural transformation with CRISPR/Cas9 counterselection. In two temporally distinct steps, we first performed a genome edit by natural transformation and second, induced CRISPR/Cas9, targeting the wild type sequence, leading to death of non-edited cells. Through highly efficient cell killing with efficiencies of up to 99.999 %, integration of antibiotic resistance markers became dispensable and thus enabled scarless and markerless edits with single-base precision. We used NT-CRISPR for deletions, integrations and single-base modifications with editing efficiencies of up to 100 % and further demonstrated its applicability for the simultaneous deletion of multiple chromosomal regions. Lastly, we demonstrated that the near PAM-less Cas9 variant SpG Cas9 is compatible with NT-CRISPR and thereby massively broadens the target spectrum.

## Introduction

*Vibrio natriegens* is a fast-growing marine bacterium with a doubling time of less than 10 minutes (Payne, 1958; Eagon, 1961) and has become a promising chassis organism due to its interesting properties, such as a wide substrate range, lack of pathogenicity and a highly active translational machinery (Aiyar, Gaal and Gourse, 2002; Weinstock *et al*., 2016; Hoffart *et al*., 2017; Hoff *et al*., 2020). A crucial factor for the utility of an organism for both basic research and application oriented projects is the availability of efficient and reliable methods for genome engineering. In case of the primary prokaryotic model organism *Escherichia coli*, large scale genome engineering projects, e.g. genome reduction (Pósfai *et al*., 2006; Kato and Hashimoto, 2007) or recoding of the *E. coli* genome by replacing all amber stop codons (Isaacs *et al*., 2011), has led to both fundamental physiological insights (Pósfai *et al*., 2006; Kato and Hashimoto, 2007) as well as to platform strains for biotechnological applications (Lee *et al*., 2009; Selas Castiñeiras *et al*., 2018). In addition to single strains with multiple modifications, a library of strains each carrying a deletion of a single non-essential gene, known as the Keio Collection (Baba *et al*., 2006), has proven to be a highly valuable resource for basic research in *E. coli*. So far, such ambitious projects with *V. natriegens* are still distant prospects and require the establishment of genome engineering methods matching those available for *E. coli* in terms of efficiency and ease of use (Wang *et al*., 2009; Jiang *et al*., 2015; Reisch and Prather, 2015).

In recent years, first steps have been made to turn *V. natriegens* into a genetically accessible model organism. Its genome, consisting of two chromosomes, was sequenced (Maida *et al*., 2013; Wang *et al*., 2013) and protocols for transformation with plasmids as well as first genome engineering methods have been developed by either taking inspiration from *E. coli* tools (Lee *et al*., 2016, 2017; Weinstock *et al*., 2016) or by harnessing organism-specific advantages of *V. natriegens*. The latter is the case for multiplex genome editing by natural transformation (MuGENT) (Dalia *et al*., 2017). Natural transformation describes the uptake of free DNA from the environment by a host encoded, multicomponent machinery and sequential integration of this DNA into the genome by RecA mediated homologous recombination (Seitz and Blokesch, 2013). This capability is widespread among *Vibrio species* and many other phylogenetically unrelated bacteria (Johnsborg, Eldholm and Håvarstein, 2007). Natural transformation was harnessed in the MuGENT method for genome engineering in *V. natriegens* by plasmid-based overproduction of the competence master regulator TfoX, thereby circumventing the need to identify the natural inducer which is chitin in many *Vibrio species* but unknown in *V. natriegens* (Dalia *et al*., 2017). While MuGENT showed a stunning efficiency with up to 10 % of the cells carrying a deletion, this method relies on the integration of an antibiotic resistance marker into a chromosomal locus for the selection of modified clones (Dalia *et al*., 2017). As a result, subsequent removal of the antibiotic resistance marker is required through additional laborious steps or the marker remains in the genome, thereby preventing future use of the respective antibiotic for selection.

A different strategy for efficiently discriminating edited and non-edited cells has been developed in a wide range of organisms through the CRISPR/Cas9 system (Reisch and Prather, 2015; Penewit *et al*., 2018; Aparicio, de Lorenzo and Martínez-García, 2019). The endonuclease Cas9 is directed to a defined genomic locus through an easily adaptable guide RNA (gRNA), thereby providing a remarkable flexibility (Jiang *et al*., 2013). Cleavage through the Cas9/gRNA complex introduces a DNA double-strand break (DSB) (Cui and Bikard, 2016) which is lethal to most prokaryotic bacteria due to the absence of the non-homologous end joining (NHEJ) pathway (Shuman and Glickman, 2007), a mechanism ubiquitous in eukaryotes (Lieber, 2010). If CRISPR/Cas9 is designed to target the wild type sequence, non-edited cells can be efficiently killed, while modified cells are immune to Cas9 cleavage and survive. When applied following the editing method, CRISPR/Cas9-based counter selection can greatly increase the fraction of modified cells and thereby the apparent editing efficiency (Reisch and Prather, 2015; Penewit *et al*., 2018; Aparicio, de Lorenzo and Martínez-García, 2019). Cell death of *V. natriegens* as a result of Cas9-mediated DNA cleavage and consequently the absence of NHEJ has been confirmed recently (Lee *et al*., 2019).

Here, we report the NT-CRISPR method combining the highly efficient genome editing through natural transformation with a CRISPR/Cas9-based counter selection strategy. We achieved editing efficiencies of up to 100 % for deletions, integrations and point mutations, and demonstrated multiplexed deletions of three genomic target regions. The NT-CRISPR method obviates the integration of antibiotic resistance markers and thereby allows scarless and markerless genome engineering in *V. natriegens*.

## Results and Discussion

### Design of and rationale behind NT-CRISPR

For NT-CRISPR, we envisioned a one-plasmid design to achieve a convenient and fast workflow. One necessary component is the master regulator for natural competence *tfoX*. Similar to a previous approach (Dalia *et al*., 2017), we use the *tfoX* gene from *Vibrio cholerae* under the control of the isopropyl ß-D-1-thiogalactopyranoside (IPTG) inducible P_tac_ promoter. Additionally, our design requires the components of the CRISPR/Cas9 system, namely *cas9* and gRNA, both driven by the Anhydrotetracycline (ATc) inducible P_tet_ promoter (Figure 1A). In our construct, we used improved variants of P_tac_ and P_tet_ (Meyer *et al*., 2019). We found that simultaneously maintaining the coding sequences for Cas9 and a gRNA targeting a genomic sequence to be a major challenge in *V. natriegens*, as even trace production of Cas9 and gRNA through leaky promoter activity were probably sufficient to introduce DNA double-strand breaks and consequently cell death. This problem persisted even when a weak ribosome binding site was used and the strongest available SsrA-derived protein degradation tag (M0050) (Stukenberg *et al*., 2021) was fused to the *cas9* sequence to reduce cellular abundance of Cas9. This strategy was previously shown to allow simultaneous maintaining the coding sequences for Cas9 and gRNA in *E. coli* (Reisch and Prather, 2015). When *V. natriegens* cells were transformed with the plasmid carrying all CRISPR/Cas9 components, we obtained only very few colonies carrying a wide range of deleterious mutations in the plasmid, rendering the CRISPR/Cas9 system non-functional. In contrast, the same plasmid with a non-binding control gRNA could be easily introduced to this host (data not shown).

**Figure 1:**
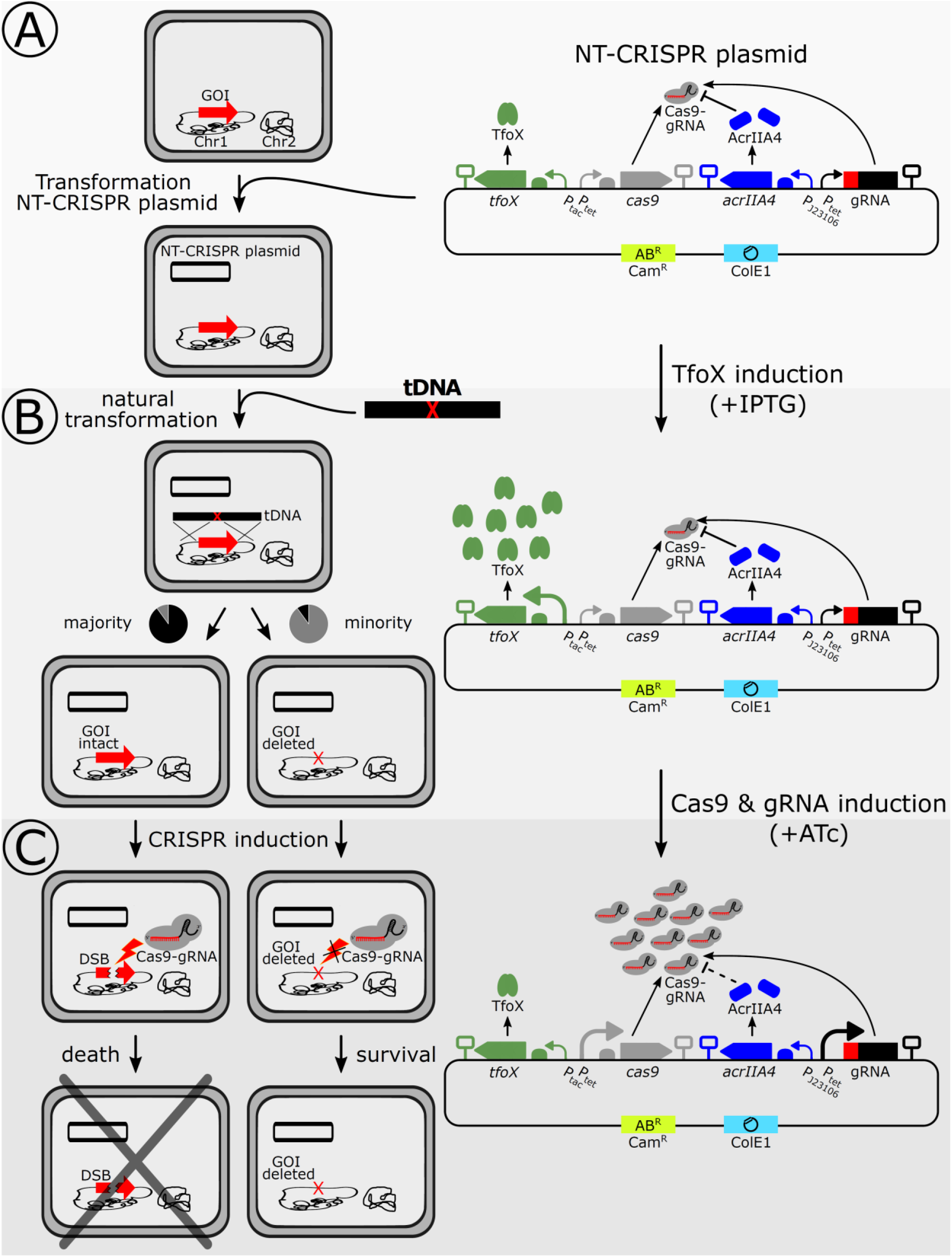
Overview of the NT-CRISPR workflow. NT-CRISPR plasmid carries tfox (P_tac_, green), cas9 (P_tet_, grey), acrIIA4 (J23106, blue) and gRNA, consisting of gRNA spacer (red) and scaffold (black), controlled by P_tet_. The backbone carries a chloramphenicol resistance marker (Cam^R^) and a ColE1 origin of replication. Increasing abundance and size of promoter indicate increased expression. (A) V. natriegens cells are tranformed with the NT-CRISPR plasmid. The gene of interest (GOI) is indicated with a red arrow on chromosome 1 (black). (B) Expression of tfox is induced by IPTG and tDNA is introduced. The tDNA is homologous to the sequences flanking the GOI. The majority of cells are not modified and the GOI stays intact, while a minority looses the GOI (red X). (C) Production of Cas9 and gRNA is induced by ATc. A DSB is introduced in wild type cells (discontinued red arrow) by Cas9-gRNA, leading to cell death. No DSB is introduced in genome edited cells, permitting survival.

The final solution to the strong toxicity of Cas9 and gRNA produced at basal levels was the expression of the anti-CRISPR protein-encoding gene *acrIIA4* under control of the constitutive promoter J23106. AcrIIA4 shows a high affinity to Cas9 and the Cas9-gRNA complex, efficiently inhibiting Cas9-gRNA from binding to the DNA target and consequently preventing the introduction of DSB (Yang and Patel, 2017; Kim *et al*., 2018). Anti-CRISPR proteins have been used in eukaryotes to achieve spatiotemporal control over CRISPR/Cas9 activity, e.g. through an optogenetic approach (Bubeck *et al*., 2018), by selective expression in certain cell types (Hoffmann *et al*., 2019) or to reduce off-target effects (Shin *et al*., 2017; Aschenbrenner *et al*., 2020) and also in the prokaryotic species *Clostridium acetobutylicum* to allow cells to carry *cas9* and *gRNA* simultaneously without toxic effects (Wasels *et al*., 2020). In a similar fashion, we used constitutive expression of *acrIIA4* to compensate for the basal production of Cas9 and gRNA. After adding *acrIIA4* to our design, we were finally able to transform *V. natriegens* with a plasmid carrying both *cas9* and a gene coding for a chromosome targeting gRNA. The full plasmid design is shown in Figure 1A.

The first step after transformation of the NT-CRISPR plasmid (Figure 1A) is the natural transformation, which requires the addition of IPTG to induce production of the master regulator TfoX. For a simple deletion, a transforming DNA (tDNA) with sequences homologous to the regions flanking the target sequence to be deleted is added to the cells (Figure 1B). A minority of cells in the population will be modified, while the vast majority of cells will still have the wild type sequence. The last step in the workflow is the CRISPR/Cas9 mediated counterselection (Figure 1C). Therefore, ATc is added to induce expression of both *cas9* and the gRNA gene to overcome the AcrIIA4 threshold causing DSB and cell death in cells with the wild type sequence. Modified cells are immune to CRISPR/Cas9 counterselection because the target sequence is no longer present in the genome (Figure 1C). As a result, the counterselection step enriches for successfully edited cells.

### Inducible cell killing by CRISPR/Cas9

As described above, it is crucial for the NT-CRISPR method to tightly control CRISPR/Cas9 and induce cell death only upon induction. We reasoned that the ratio between *acrIIA4* and *cas9* expression strength is important. Too weak *acrIIA4* expression might not be enough to compensate for the leaky *cas9* and *gRNA* gene expression and still lead to premature cell death, while too strong *acrIIA4* expression could prevent efficient counterselection even with a full induction of the CRISPR/Cas9 system. To test a range of *acrIIA4* and *cas9* expression strengths, we created twelve plasmids representing the combinatorics of four and three different ribosome binding sites (RBS) for *acrIIA4* and *cas9*, respectively. The RBS used differ in their expression strength which was recently quantified in *V. natriegens* with fluorescent reporter experiments (Stukenberg *et al*., 2021). These variants were tested for their ability to trigger cell death upon addition of ATc as the inducer of Cas9 and gRNA production using a gRNA targeting the non-essential *wbfF* gene, which is involved in capsule polysaccharide biosynthesis (Bik *et al*., 1996). Growth of cells cultivated in microplates was continuously measured and ATc was added in saturating concentrations (200 ng/mL) to exponentially growing cells 1h after start of the batch cultivation. For all strains carrying plasmids using the moderately strong RBS B0032 (Stukenberg *et al*., 2021) for *acrIIA4*, we observed a reduction in OD_600_, indicating cell death, about 1.5 h after induction of *cas9* and the gRNA with ATc (Supplementary Figure S1A). While the choice of the RBS for *acrIIA4* and consequently its expression strength, apparently has a strong impact on the inducibility of the CRISPR/Cas9 system, we found no difference between the three tested RBS for *cas9*, neither in this growth experiment in liquid culture (Supplementary Figure S1A) nor in a separate agar plate-based assay which resembles the counterselection efficiency within the actual NT-CRISPR editing workflow (Supplementary Figure S1B). Of the three variants, which showed inducible cell killing in the growth experiment, we proceeded with the plasmid pST_116 which carries the strongest of the three tested RBS for *cas9* B0033 (Stukenberg *et al*., 2021).

### Dependence of gene deletion efficiencies on tDNA amount and homologous sequence length

As a first target to demonstrate gene deletion by our NT-CRISPR-based approach in *V. natriegens*, we again chose *wbfF*, since deletion mutants in this gene show an almost transparent colony morphology (Dalia *et al*., 2017). This phenotype allows for distinction between wild type and *wbfF* mutant colonies to conveniently calculate gene deletion efficiencies. We reproduced this phenotype by our approach using tDNA comprising 3 kb homologous sequences flanking the designed deletion (Figure 2A). Subsequently, we characterized the genome editing efficiency dependent on the amount of tDNA supplied (1 ng, 10 ng or 100 ng) and on the induction state of the CRISPR/Cas9 system (Figure 2B). We obtained close to 100 % of all colonies with the transparent Δ*wbfF* morphology for all three DNA amounts tested when CRISPR/Cas9 was induced. This suggests a remarkably high editing efficiency of NT-CRISPR, even when just 1 ng of tDNA was used. However, in absence of counterselection, we obtained a lower fraction of positive colonies which decreased with reduction in tDNA amounts (Figure 2B). This is in accordance with a previous study describing natural transformation for genome engineering in *V. natriegens* (Dalia *et al*., 2017). This trend also became apparent through the absolute number of positive colonies (CFU/µL) which for both the induced and uninduced samples decreased with lower amounts of tDNA used (Supplementary Figure S2A, S2B). With 100 ng of added tDNA and without counterselection, we still obtained ∼ 20 % of positive colonies (Figure 2B), further supporting the potential of natural transformation for genome engineering in *V. natriegens*.

**Figure 2:**
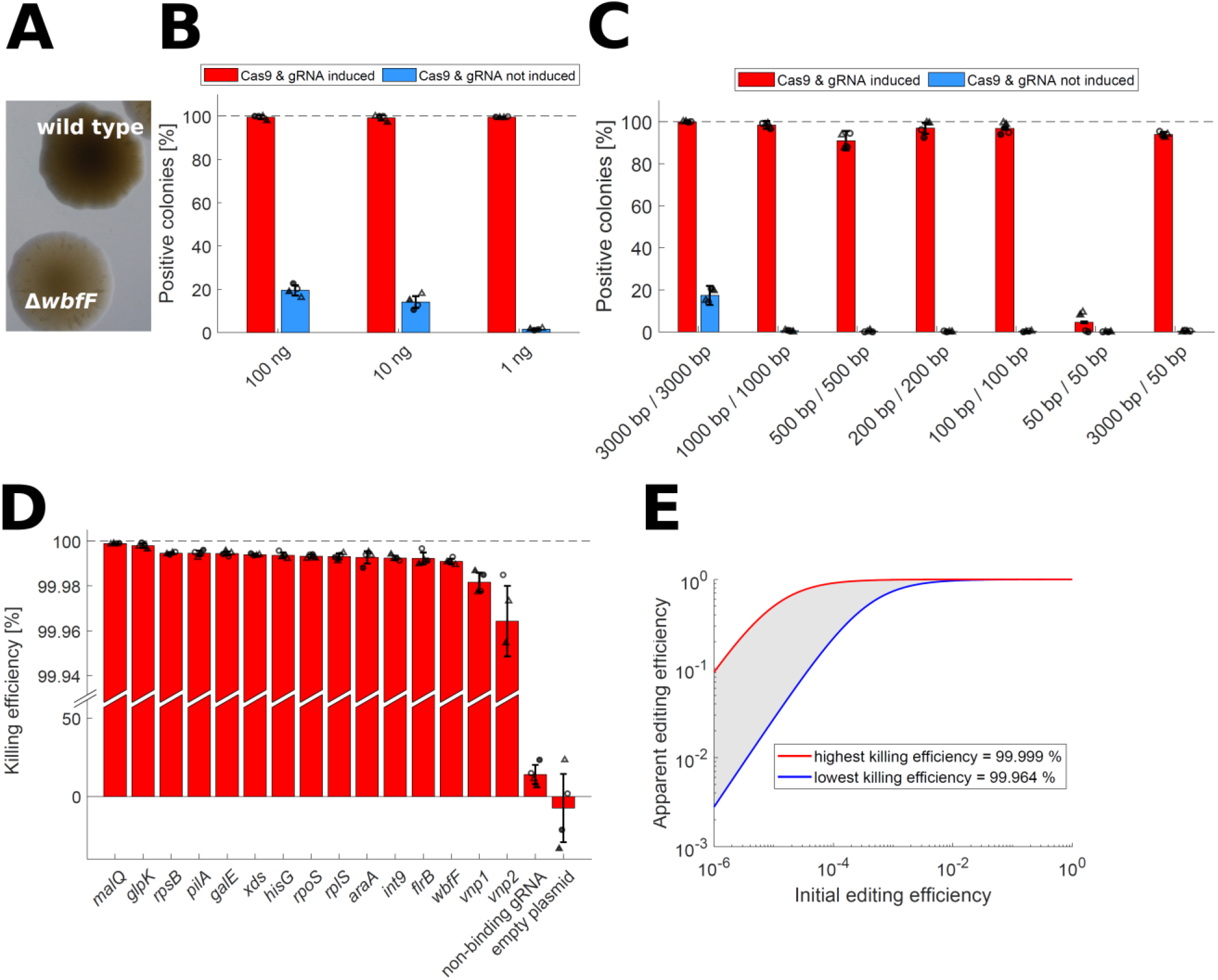
Quantitative characterization and killing efficiencies. (A) Different morphologies of wild type and ΔwbfF colonies. Image acquired with transmission light. (B) NT-CRISPR with different amounts of tDNA, targeting wbfF. This experiment was performed with the indicated amount of tDNA with symmetrical 3 kb flanks upstream and downstream of the target sequence. Red and blue bars show results for samples with and without CRISPR-based counterselection, respectively. Data represent the mean of two biological replicates (circle or triangle) and two independent experiments (filled or open symbols). Error bars indicate standard deviation of the mean. Underlying colony counts are provided in Supplementary Figures S2A and S2B. Dashed line indicates the highest possible value. (C) NT-CRISPR with different tDNA fragment lenghts, targeting wbfF. Red and blue bars show results for samples with and without CRISPR-based counterselection, respectively. This experiment was performed with 100 ng of tDNA and the respective fragment length. With the exception of the last bar, tDNA fragments are symmetrical with the same length for the upstream and downstream homologous sequence. Data represent the mean of two biological replicates (circle or triangle) and two independent experiments (filled or open symbols). Error bars indicate standard deviation of the mean. Underlying colony counts are provided in Supplementary Figures S2C and S2D. Dashed line indicates the highest possible value (D) Results of killing assay for different gRNAs. Killing efficiency is calculated as follows: 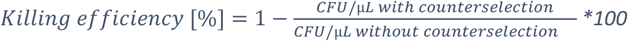. Data represent the mean of two biological replicates (circle or triangle) and two independent experiments (filled or open symbols). Error bars indicate standard deviation of the mean. Dashed line indicates the highest possible value. (E) Calculation of apparent editing efficiency from initial editing efficieny. Initial editing efficiency describes a theoretical value, assuming no enrichment through counterselection against non-modified cells. Apparent editing efficiency provides the expected fraction of correct colonies after counterselection. The red and blue curves use the highest (malQ) and lowest (vnp2) killing efficiencies observed in Figure 2D.

In a second experiment, we investigated the impact of different lengths of the homologous sequences of the tDNA on the success of genome editing with NT-CRISPR (Figure 2C). When CRISPR/Cas9 was induced, we again obtained high editing efficiencies of at least 90 % for all tested fragment lengths, with exception of the samples with 50 bp homologous flanks. More than 98 % editing efficiency was achieved for 3000 bp and 1000 bp homologous flanks. However, without counterselection, we obtained less positive colonies and also the CFU/µL with Δ*wbfF* mutant morphology declined with decreasing length of the homologous flanks (Supplementary Figure S2C, S2D). Decreases in putative Δ*wbfF* CFU/µl by ∼ 40 fold and ∼ 4 fold were observed when reducing the flanks from 3000 bp to 1000 bp and 1000 bp to 500 bp, respectively (Supplementary Figure S2C). Further shortening the flanks down to 100 bp did not result in considerable additional drops in putative Δ*wbfF* CFU/µl (Supplementary Figure S2C). Note that using the same mass of tDNAs of different lengths equates to more molecules of the shorter tDNA fragments, suggesting that a possible reduction of uptake or recombination efficiency of shorter fragments is partly compensated by a higher concentration of tDNA molecules. Another result with potential practical implications is the high editing efficiency of ∼ 90 % obtained by NT-CRISPR with an asymmetric tDNA fragment with 3000 bp and 50 bp homologous sequence surrounding the deletion (Supplementary Figure S2C, S2D), as this fragment can be easily generated in one PCR by adding the short 50 bp arm as an overhang to one of the PCR primers used to amplify the long arm. Applying such an asymmetric DNA fragment for natural transformation-mediated locus exchange by homologous recombination was successfully established for *V. cholerae*, but required the deletion of at least two ssDNA exonucleases to achieve reasonable editing efficiencies (Dalia *et al*., 2017). If deletion of the orthologous ssDNA exonucleases in *V. natriegens* could increase the editing efficiency with short tDNA fragments even further remains unknown. Nonetheless, due to the highly efficient counterselection, asymmetrical tDNA fragments with one short arm might represent a convenient approach to introduce single gene deletions.

Collectively, the initial characterization suggests that combining natural transformation with efficient, targeted CRISPR Cas9/gRNA-mediated counterselection considerably increases genome editing frequencies in *V. natriegens* and that even low amounts of tDNA like 1 ng and homologous flanks as short as 100 bp are sufficient to produce more than 90 % correct colonies.

### NT-CRISPR proof-of-concept for generating deletions of different lengths, point mutations and integrations in a set of target genes

Inspired by these promising results, we sought to apply CRISPR-NT for various types of genome edits and to expand the set of chromosomal target sequences for our proof-of-concept study in *V. natriegens*. We assembled the NT-CRISPR plasmids with 15 different gRNAs, targeting a range of different sequences, both on chromosome 1 and 2. The reason behind selecting these genes is described below. In order to quantify the “killing efficiency” of the respective gRNAs, we performed the NT-CRISPR protocol without addition of tDNA. Samples were either induced with ATc to induce CRISPR/Cas9 counterselection or remained uninduced. The ratios of the resulting CFU/µL allows calculation of killing efficiency in the NT-CRISPR workflow. The tested gRNAs show killing efficiencies between 99.999 % (*malQ*) and 99.964 % (*vnp1*) (Figure 2D). The value of the strongest tested gRNA that targeted *malQ* translates to the survival of just one out of 100.000 cells. A non-targeting gRNA was included as a control and resulted in a slight reduction of cells upon Cas9 and gRNA induction, possibly due to unspecific toxicity or protein expression burden. An empty plasmid control ruled out negative effects of the inducer ATc at the applied concentration.

Knowing the killing efficiency, it is possible to calculate the apparent editing efficiency, defined as the fraction of positive cells after counterselection, from the initial editing efficiency before counterselection. With a highly efficient gRNA even just 10^−6^ (one edited cell out of one million) results in 10^−1^ or 10 % of all remaining cells being correct (Figure 2E). However, even with the weakest tested gRNA, an initial editing efficiency of 10^−3^ (0.1 %), which is far below the values obtained for simple deletions without counterselection (Figure 2B), is sufficient to achieve apparent editing efficiencies of almost 100 % through robust counterselection.

#### Deletions

To kill two birds with one stone, we picked genes and groups of genes as targets for deletion which knockout might lead to increased plasmid transformation efficiencies. So far, the tremendous potential of *V. natriegens* as a fast-growing strain for the selection and propagation of *in vitro* assembled plasmids is still hampered due to transformation efficiencies that are much lower than the ones observed for highly engineered *E. coli* strains (Stukenberg *et al*., 2021). We targeted the two prophage regions *vnp1* and *vnp2*, as their removal leads to an increased robustness towards osmotic stress (Pfeifer *et al*., 2019) which might be experienced by the cells during the preparation of chemically competent cells. Additionally, we chose *galE*, an enzyme which provides precursors for the synthesis of lipopolysaccharides because an increased plasmid transformation efficiency was reported for *galE* mutants of some Gram-negative bacteria, presumably due to the loss of cell surface structures which might impair plasmid uptake (Bursztyn *et al*., 1975; Van Die *et al*., 1984). Along the same line, we reasoned that removal of other surface structures, namely the flagella and pili, could also contribute to increased transformation efficiency. Lastly, we included the gene coding for the extracellular nuclease Xds (Blokesch and Schoolnik, 2008), which could degrade the incoming plasmid DNA similarly to the Dns nuclease, as its deletion was already confirmed to drastically increase the plasmid transformation efficiency of chemically competent cells (Weinstock *et al*., 2016; Stukenberg *et al*., 2021).

The targets for deletion described above span a wide range in size from 1.0 kb (*galE*) up to the prophage region of *vnp2* with 39 kb and are located on either chromosome 1 or chromosome 2 (Figure 3A). To quantify the editing efficiency of these deletions, we analyzed 50 colonies by colony PCR. For all targets, except for the two prophage regions, all screened colonies were correct, including the deletion of a flagellar gene cluster of 31 kb. Also the prophage regions were deleted with 47 and 49 colonies out of 50 being correct, a remarkably high efficiency considering the size of 36 kb and 39 kb, respectively. We sequenced the target regions for four colonies per target and found the desired sequence in all colonies, with a single exception. In case of *vnp1*, one colony missed four bp directly adjacent to the deleted sequence (Supplementary Figure S3). It was shown previously, that the prophages are activated spontaneously at low frequencies (Pfeifer *et al*., 2019). It is tempting to speculate that this deviation from the desired sequence is the result of a spontaneous loss of the prophage region, which would still confer resistance to the CRISPR-based counterselection, instead of the targeted deletion by natural transformation.

**Figure 3:**
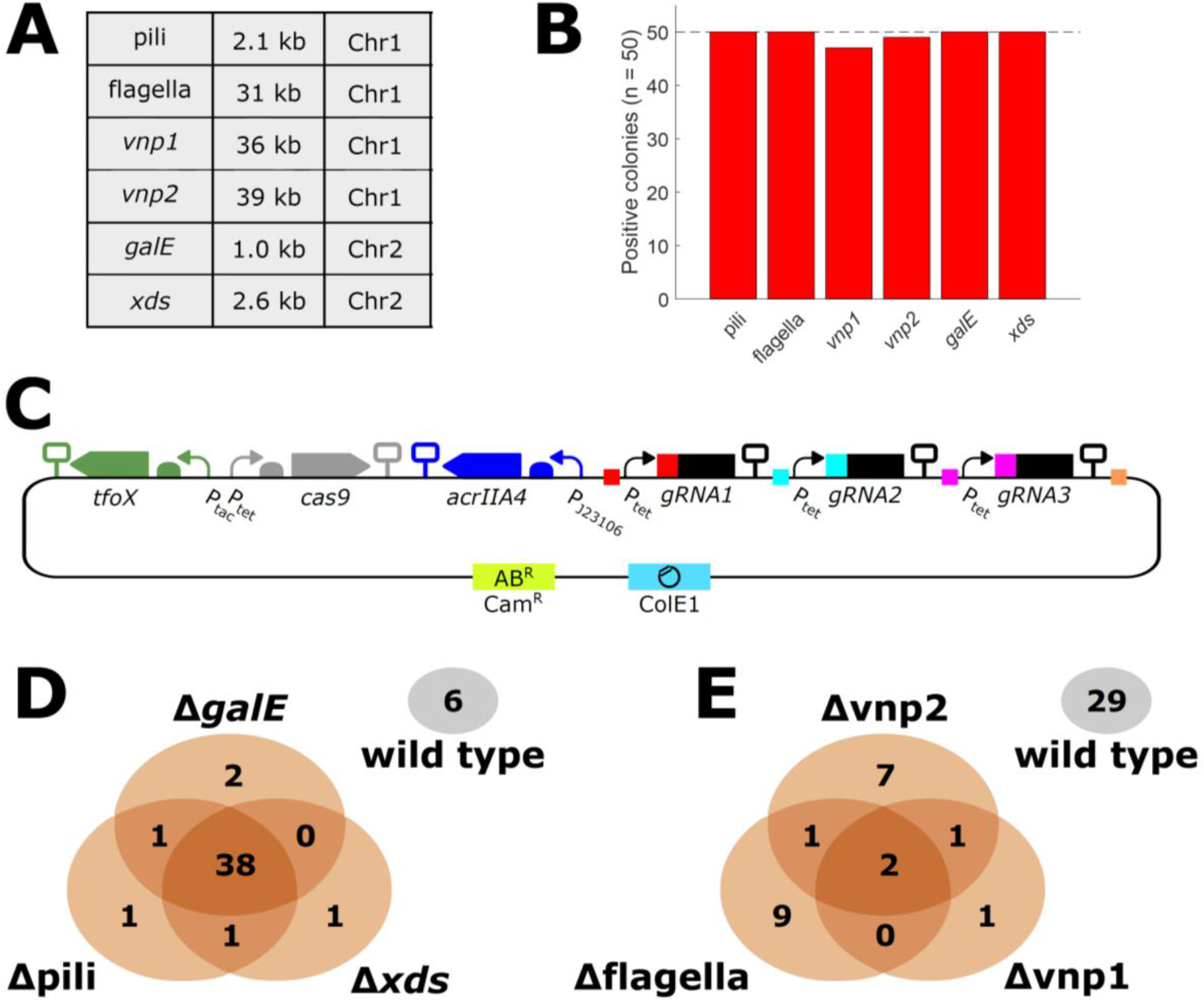
Results of deletions with NT-CRISPR. (A) Table providing information about deleted sequence. Locus tags of deleted genes are as follows: Pili (PN96_01310 - PN96_01315), flagella (PN96_02540 - PN96_02685), vnp1 (PN96_04290 - PN96_04520), vnp2 (PN96_06880 - PN96_07085), galE (PN96_22140), xds (PN96_19285). Chr1 = Chromosome 1, Chr2 = Chromosome 2. (B) Efficiency of deletions. Results obtained by PCR (n = 50). Dashed line indicates the highest possible value. (C) Visualization of NT-CRISPR plasmid carrying three gRNAs. Colored squares indicate matching fusion sites used for construction of this plasmid using Golden Gate Assembly. More details regarding the assembly of a multi gRNA NT-CRISPR plasmid is provided in Supplementary Figure S5 (D, E) Venn diagrams to visualize results of multigene deletions. Note that areas of ellipses and intersections are not proportional to the displayed values. Colonies carrying none of the deletions are indicated with a separate ellipsis (grey). Results obtained through PCR (n = 50). Plasmids used are pST_138 (D) and pST_137 (E).

As described above, the targets for deletion were selected because we hoped that their deletion would improve plasmid transformation efficiency of *V. natriegens*. We tested all single deletions but did not see any significant increase (Supplementary Figure S4).

One feature of CRISPR-based systems is its inherent modularity. Simultaneous targeting of multiple sequences is possible by simply including multiple gRNAs (Figure 3C). Assembly of a plasmid harboring all required components to target three different loci is achieved by firstly constructing the gRNA expression cassette and, secondly, by integrating them into a plasmid already carrying all remaining components (Supplementary Figure S5). We tested this approach by simultaneously deleting three of the genomic regions which could be efficiently removed individually. We obtained striking results for the simultaneous deletion of the pili operon, *xds* and *galE*, with 38 out of 50 tested colonies carrying all three deletions (Figure 3D). As an even bigger challenge, we successfully deleted the flagellar gene cluster and both prophage regions simultaneously in two out of 50 colonies (Figure 3E). The deleted regions sum up to 106 kb, equaling ∼ 2 % of the *V. natriegens* genome. Killing efficiency with three gRNAs was found to be similar or slightly higher than that with the individual gRNAs (Supplementary Figure S6), suggesting that undesired recombination events within the sequences identical in all gRNAs (P_tet_ and gRNA scaffold) did not occur at perturbing frequencies.

#### Point mutations

In addition to deletions, we tested NT-CRISPR for the introduction of point mutations. Three genes, *malQ, araA* and *glpK*, involved in catabolism of the alternative carbon sources maltose, arabinose and glycerol, respectively, were chosen as proof-of-concept targets for the introduction of point mutations because introduction of a premature stop codon by a single point mutation can be easily identified through loss of *V. natriegen’s* ability to grow on minimal medium with the respective carbon source. In case of *malQ*, we randomly selected 50 colonies yielded by the NT-CRISPR approach and tested them for their ability to grow on M9 minimal medium with maltose as the sole carbon source. None of the tested colonies could grow indicating successful genome modification (Figure 4A). Subsequently, the target sequence of four colonies was sequence and the introduction of the desired point mutation was confirmed for all tested colonies. In case of *araA* and *glpK*, a first attempt to introduce a point mutation was not successful and resulted in very few colonies, none of them carrying the desired edit (data not shown). High editing efficiency of 100 %, based on the colony’s inability to grow on minimal medium with the respective carbon source, could be achieved by introducing a second silent point mutation, leading to a C-C mismatch (G->C mutation) (Figure 4A). It was shown previously that *V. natriegens* has an active methyl-directed mismatch repair (MMR) pathway, preventing the efficient introduction of point mutations by natural transformation (Dalia *et al*., 2017). It is known that C-C mismatches inhibit MMR in a wide range of other organisms (Su *et al*., 1988; Detloff, Sieber and Petes, 1991; Meier and Wackernagel, 2005) and our data suggest that this is also the case in *V. natriegens*. The approach to introduce a C-C mismatch in addition to the desired mutation can serve as a convenient solution, when MMR hinders successful introduction of certain point mutations. Again, we confirmed the point mutations, as well as the additional C-C mismatch by sequencing of the target regions of *glpK* and *araA*.

**Figure 4:**
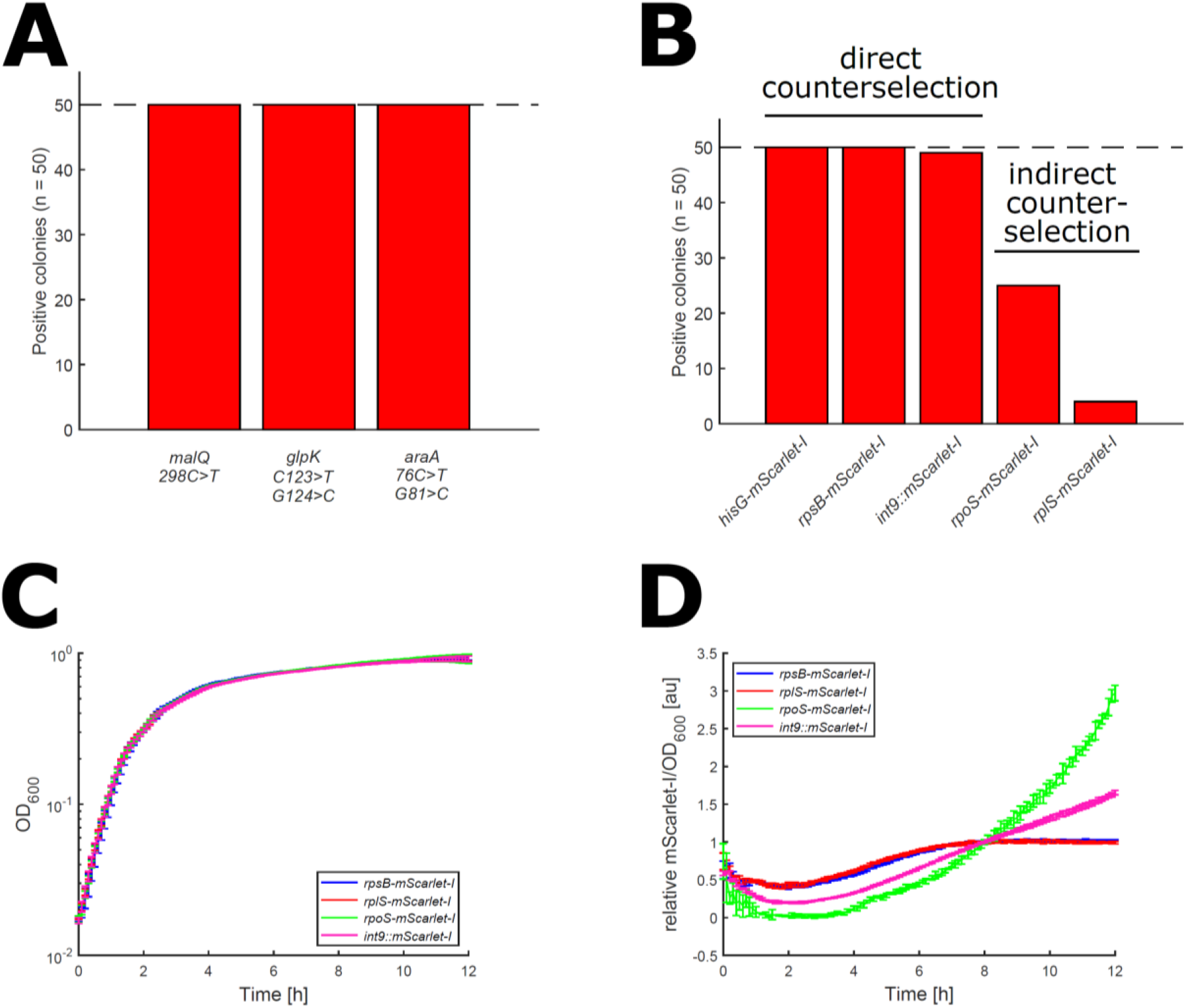
Results of point mutations and integrations. (A) Efficiency of introducing point mutations. Targeted genes are malQ (PN96_15600, Chr1), glpK (PN96_01955, Chr1) and araA (PN96_16040, Chr2). Results obtained through ability to grow on the respective carbon source. First indicated mutation introduces a stop codon and second mutation (for glpK and araA) introduce a silent mutation as a C-C mismatch to evade mismatch repair. (B) Integration of mScarlet-I. mScarlet-I fused to 3’ end of hisG (PN96_07800), rpsB (PN96_02260), rpoS (PN96_01115) and rplS (PN96_01280) and the integration of mScarlet-I with a constitutive promoter into one intergenic regionbetween genes with locus tags PN96_06135 and PN96_06140, all on chromosome 1. “Direct counterselection” describes counterselection through a gRNA overlapping the integration site, while “indirect counterselection” refers to the selection through a silent point mutation ∼ 300 bp upstream of the integration site. (C) Growth curves of translational fusions and strain with integrated constitutive mScarlet-I cassette. Data represent the mean of four biological replicates and two independent experiments. Error bars indicate standard deviation from the mean. (D) Normalized mScarlet-I signal. mScarlet-I/OD_600_ was normalized by value at timepoint 8 h to compensate for different mScarlet-I signals. Underlying data without normalization is shown in Supplementary Figure S7. Data represent the mean of four biological replicates and two independent experiments. Error bars indicate standard deviation from the mean.

#### Integrations

Lastly, we wanted to test if NT-CRISPR is also applicable for integrations into the genome. The approach presented here could be a powerful tool to fuse fluorescent reporter genes to any gene of interest to study their expression or localization of their gene product. We selected genes which expression is expected to differ during the course of growth in a batch culture to follow this dynamic by measuring the reporter signal. We fused the coding region of the red fluorescent protein mScarlet-I (Bindels *et al*., 2017) to the 3’ end of four genes which are known to be differentially regulated during transition into and in the stationary phase in *E. coli*. As candidates for genes with high expression level in the exponential growth phase, we chose two ribosomal protein encoding genes. We picked *rpsB* and *rplS* because fusion of fluorescent reporter proteins to the respective ribosomal proteins was possible in *E. coli* without noticeable detrimental effects on growth or ribosome assembly (Nikolay *et al*., 2014). As candidates for genes upregulated in stationary phase, we chose *hisG*, involved in histidine biosynthesis (Cavanagh, Chandrangsu and Wassarman, 2010) and *rpoS* encoding the stress sigma factor σ^38^ (Jaishankar and Srivastava, 2017). A control strain was generated by equipping an mScarlet-I transcription unit with a strong constitutive promoter (J23111) and a strong RBS (B0030), and integrating this construct into one intergenic region with neighboring genes in convergent orientation (int9).

Counterselection for successful integration can be performed through a gRNA which overlaps the integration site so that successful modification disrupts the gRNA binding sequence and thereby confers resistance against CRISPR-based counterselection. In case of *rpoS* and *rplS*, no suitable PAM sequence was available at the desired integration site. As a workaround, we introduced a silent point mutation 300 bp from the integration site and used this point mutation for the counterselection, expecting the integration of *mScarlet-I* when the silent point mutation is present. For each integration, 50 randomly selected colonies were screened by colony PCR. When a gRNA was available for direct counterselection at the integration site, we reliably obtained high editing efficiencies with almost all tested colonies being correct (Figure 4B). In contrast, for *rpoS* and *rplS* editing efficiencies were drastically lower with just 25 and 4 colonies, respectively, carrying the desired *mScarlet-I* integration. Sequencing of four colonies each, which were negative in the PCR screening, revealed that all carried the selected silent point mutation, suggesting that not the full-length tDNA fragment was incorporated through homologous recombination. It remains to be investigated if introduction of the silent point mutation closer to the actual integration site than the 300 bp tested here, might lead to better results. The generated constructs were tested for their growth behavior and mScarlet-I signal. No growth difference was observed between all tested strains (Figure 4C), including *mScarlet-I* fusions to the essential ribosomal genes *rpsB* and *rplS* (Lee *et al*., 2019) which is in accordance with similar experiments in *E. coli* (Nikolay *et al*., 2014). The mScarlet-I signal of the tested strains showed the expected growth phase dependency. The *rpsB-mScarlet-I* and *rplS-mScarlet-I* strains showed an identical pattern with a steady increase in reporter-mediated fluorescence signal till 6 h, before reaching a plateau. In contrast, the *rpoS-mScarlet-I* strain showed an increasing signal after 4 h with an even steeper rise after 6h. A constant increase from the onset of the stationary phase (∼ 2 h) and throughout the stationary phase was observed for the strain carrying the constitutively expressed *mScarlet-I* cassette (Figure 4D). The data are shown as relative mScarlet-I/OD_600_ to account for strong differences in the absolute signal strength (Supplementary Figure S7). Unfortunately, the signal for *hisG-mScarlet-I* was hardly measurable and did not allow for quantitative characterization (data not shown). The results presented here serve as an example how NT-CRISPR can be used for the construction of reporter strains to study the regulation of important physiological processes.

### Overcoming PAM dependency with engineered Cas9

The use of Cas9 is limited by the availability of the PAM sequence, NGG (N = any nucleotide) in case of the commonly used Cas9 from *Streptococcus pyogenes*. While there are always plenty of possible gRNAs available for larger deletions, introduction of specific point mutations or genomic integrations into a desired locus might be restricted when no PAM can be found nearby, thus requiring inefficient workaround solutions (see generation of *rpoS-mScarlet-I* and *rplS-mScarlet-I* above). In recent years, huge progress has been made in developing new Cas9 variants with a wider PAM spectrum (Kleinstiver *et al*., 2015; Hu *et al*., 2018; Walton *et al*., 2020). We tested the near-PAMless Cas9 variant SpG Cas9 (Walton *et al*., 2020) with the PAM requirement being NGN. In theory, each G and C nucleotide in the genome can serve as a PAM, thereby massively expanding the number of available gRNAs.

We introduced the described mutations into the *cas9* sequence in the NT-CRISPR plasmid and tested it with gRNAs targeting *wbfF* with all possible PAM doublet pairs (NGG, NGA, NGC, NGT). The *wbfF* gRNA with NGG PAM showed similar killing efficiencies when used with Cas9 and SpG Cas9, confirming the general compatibility of SpG Cas9 with NT-CRISPR. The tested gRNAs with alternative PAM sequences showed a wide range of killing efficiencies from 99.993 % (NGT) and no significant effect for (NGC) (Figure 5A). We note that other determinants, apart from the PAM sequence, can influence the killing efficiency and the limited number of tested gRNAs does not allow formulation of general claims about the applicability of SpG Cas9 with alternative PAMs in the framework of NT-CRISPR in *V. natriegens*. However, it appears as if the killing efficiencies were far lower than the obtained values for the many gRNAs tested before with NGG PAMs in our study (cf. Figure 2D), except for the tested gRNA using NGT as PAM. Based on the high killing efficiency which we observed with a gRNA using NGT as PAM, we designed two additional gRNAs overlapping the 3’ end of *rpoS* and *rplS* and also measured high killing efficiencies for these (Figure 5A). Thereafter, we used SpG Cas9 together with these two gRNAs for the integration of *mScarlet-I* at the 3’ end of *rpoS* and *rplS*. For 50 randomly selected colonies, successful mScarlet-I integration was confirmed for 44 and 50 colonies for rpoS and rplS, respectively, compared to 25 and 4 for the indirect approach described above (Figure 5B). In conclusion, the killing efficiency with SpG Cas9 and alternative PAM sequences tends to be lower than Cas9 with NGG PAMs, but our results suggest that applying SpG Cas9 together with gRNAs using NGT as PAM can be a suitable strategy for the integration of sequences when no gRNA with NGG is available at the desired target sequence.

**Figure 5:**
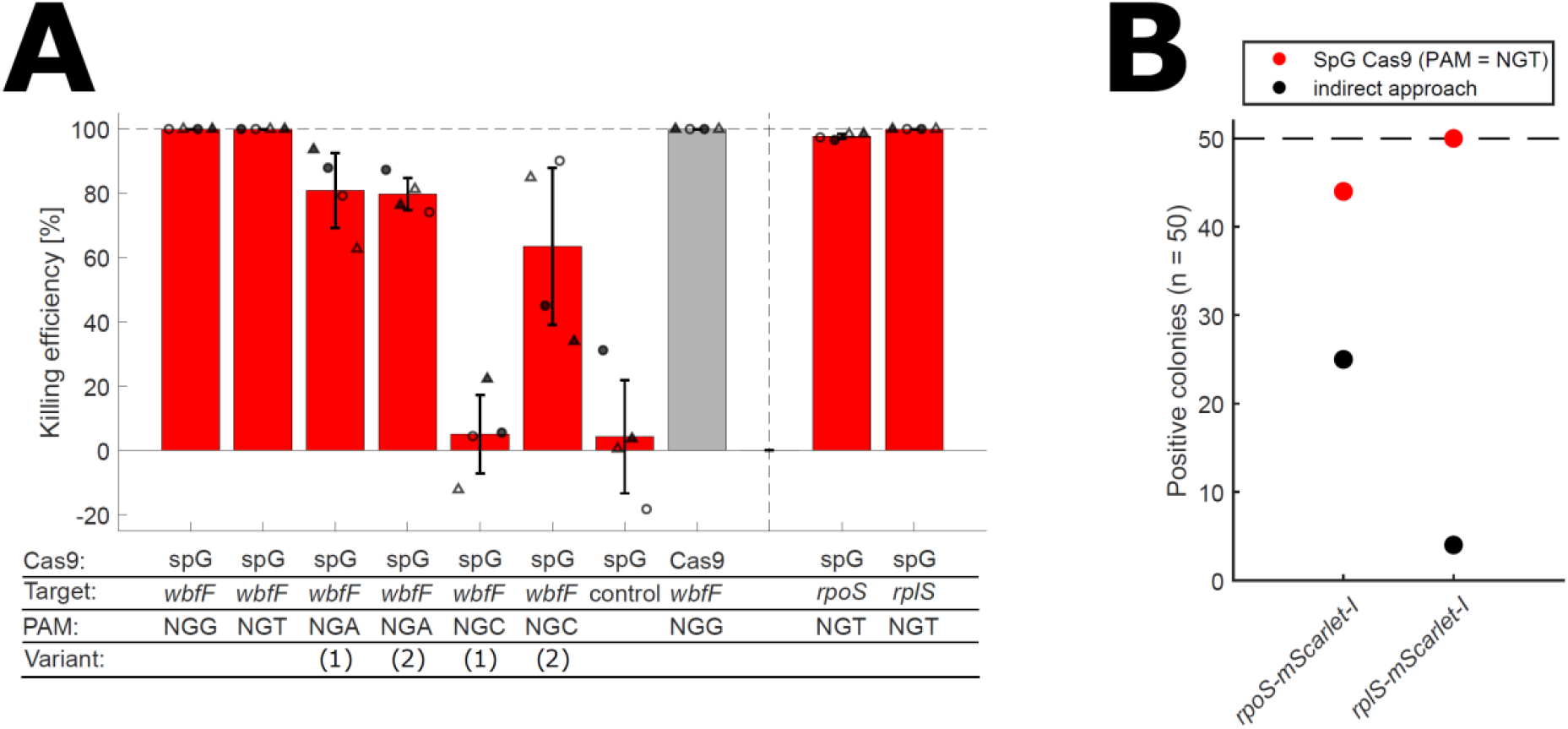
(A) Testing spG Cas9 in NT-CRISPR. Killing efficiency with spG Cas9 with all possible PAM sequences. Cas9 with NGG PAM is shown as a reference in grey. Killing efficiency is calculated as follows: 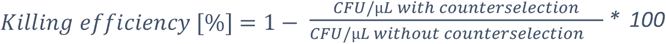. Data represent the mean of two biological replicates (circle or triangle) and two independent experiments (filled or open symbols). Error bars indicate standard deviation of the mean. Dashed line indicates the highest possible value. (B) Efficiency for integration of mScarlet-I using either SpG Cas9 with a NGT PAM sequence or using the indirect approach described above.

## Conclusion

In this study, we developed NT-CRISPR which builds on the previous application of natural transformation for genome engineering in *V. natriegens* (Dalia *et al*., 2017) by adding a CRISPR/Cas9 counterselection step. With killing efficiencies of up to 99.999 %, most genomic modifications, including deletions, integrations, and point mutations, can be performed with almost 100 % efficiency. As one highlight, we demonstrate the simultaneous deletion of multiple genes through the expression of multiple gRNAs directed against different target sequences. The NT-CRISPR workflow can be performed in a standard working day and the full process, including preparation of tDNAs and cloning of gRNAs, as well as curing of the NT-CRISPR plasmid after successful genome modification, can be achieved in one week. Edits with this method can be performed with single base precision, without the – transient or permanent – integration of antibiotic resistance markers and do not leave any undesired scars in the genome. The major limitation of this method is its PAM dependency which might restrict access to some sequences for modifications. We present two approaches to close this gap, by either introducing a silent, selectable point mutations or by using SpG Cas9 (Walton *et al*., 2020) with alternative PAM sequences.

We are confident that this method will provide the growing *V. natriegens* community with a convenient and highly efficient genome engineering tool. It sets the foundation for sophisticated strain engineering projects to exploit the fascinating properties of *V. natriegens* for academic and industrial applications in the future.

Furthermore, natural transformation is a commonly used strategy for genome engineering in a wide range of biotechnologically and clinically relevant bacteria, e.g. *Bacillus subtilis* (Vojcic *et al*., 2012), *Vibrio cholerae* (Dalia, McDonough and Camilli, 2014), *Vibrio fischeri* (Visick *et al*., 2018), *Streptococcus thermophiles* (Blomqvist, Steinmoen and Håvarstein, 2006) and *Streptococcus mutans*. The NT-CRISPR method described here for *V. natriegens* could serve as a blueprint to upgrade existing natural transformation approaches with CRISPR/Cas9-based counterselection to accelerate research in these important organisms.

## Methods

### Bacterial strains and culture conditions

The *V. natriegens* strain used for this study is a derivative of ATCC14048 with deletion of the *dns* gene, constructed in a previous project for increased plasmid transformation efficiency (Stukenberg *et al*., 2021). *V. natriegens* was routinely grown in LBv2 (Weinstock *et al*., 2016) medium with added chloramphenicol if appropriate. Chloramphenicol was added with a final concentration of 4 µg/mL for liquid and 2 µg/mL for solid medium.

Strains were prepared for long term storage at -80 °C by growing cultures for 6 - 8 h at 37 °C and mixing 700 µL of grown cultures with 700 µL of 50 % glycerol. We found that the additional washing step reported previously (Stukenberg *et al*., 2021) is not required when cultures are not grown into deep stationary phase (e.g. overnight at 37 °C).

### Preparation of chemically competent *V. natriegens* cells and heat-shock transformation

Preparation of chemically competent cells and heat-shock transformation was performed exactly as described before (Stukenberg *et al*., 2021). A preculture of *V. natriegens* ATCC14048 Δ*dns* was inoculated from a glycerol stock and grown overnight at 37 °C and 220 rpm. At the next day 125 mL of preheated LBv2 medium (37 °C) was inoculated with the overnight culture to a final OD_600_ of 0.01 in a 1 L baffled shake flask. This culture was grown at 200 rpm (37 °C) until an OD_600_ between 0.5 and 0.7 was reached. The culture was then transferred to pre-cooled 50 mL falcon tubes and incubated on ice for 10 min, followed by centrifugation for 10 min at 3000 x g at 4 °C. The supernatant was discarded, and the pellets were resuspended in 40 mL cold TB buffer per 125 mL bacterial culture (TB buffer: 10 mM Pipes, 15 mM CaCl_2_, 250 mM KCl, pH adjusted to 6.7 with KOH, then add 55 mM MnCl_2_, filter sterilized). The cells were again incubated on ice for 10 min and further centrifuged for 10 min at 3000 x g at 4 °C. The supernatant was removed, and pellets were resuspended in 5 mL cold TB buffer per 125 mL starting culture and consolidated in a single falcon tube, before adding 350 µL Dimethyl sulfoxide (DMSO). After another 10 min incubation on ice, 50 µL aliquots were prepared in 1.5 mL tubes and snap frozen in liquid nitrogen. Aliquots were stored at -80 °C until further use.

Chemically competent *V. natriegens* ATCC14048 Δ*dns* cells were transformed by adding DNA to an aliquot of competent cells, and incubated on ice for 30 min. After 30 min, cells were heat shocked in a water bath at 42 °C for 45 s then immediately incubated on ice for 10 min before recovery. The cells were recovered in 1 mL warm LBv2 medium, shaking at 37 °C for 1 h at 700 rpm. After recovery, the cells were pelleted by centrifugation at 3000 x g for 1 min, the supernatant was decanted, and the pellet was resuspended in the remaining ∼ 50 µL residual medium. The whole volume was plated on 37 °C warm LBv2 plates containing the appropriate antibiotic and incubated over night at 37 °C.

### Construction of NT-CRISPR plasmids

Plasmids were constructed within the framework of the Marburg Collection, a recently published Golden Gate-based cloning toolbox for *V. natriegens* (Stukenberg *et al*., 2021). Part sequences and detailed description of plasmid assembly are provided in Supplementary Table S1 and S2, respectively.. Assembly of the plasmids was performed in *E. coli* NEB Turbo, with the exception of the exchange of gRNA spacer sequences, as the resulting plasmids are intended for experiments in *V. natriegens*. Adaptation for different target sequences was achieved by replacing a sfGFP dropout fragment with the gRNA spacer sequence by annealing two complementary oligonucleotides. If not indicated otherwise, pST_116 was used for plasmids carrying single gRNAs with Cas9 and pST_140 whenever spG Cas9 was used. Annealing reaction was set up by mixing 1.5 µL of each oligonucleotide (100 fM) with 5 µL T4-DNA ligase buffer (Thermo Scientific) in a total reaction volume of 50 µL. Reactions are incubated in a heat block at 95 °C for 15 min, before switching off the heat block to allow samples to cool down slowly to room temperature (∼ 1 h). Cloning reaction with the NT-CRISPR plasmids was set up with ∼ 200 ng of the respective plasmid, 3 µL annealed oligonucleotides, 0.5 µL of T4-DNA Ligase (5 Weiss U/µL, Thermo Scientific) and BsaI (10 U/µL) and 1 µL T4-DNA ligase buffer in a total reaction volume of 10 µL. Reactions were run in a thermocycler with 30 cycles of 37 °C (2 min) and 16 °C (5 min), followed by a final digestion step at 37 °C for 30 min and an enzyme denaturation step at 80 °C for 10 min. Transformation of *V. natriegens* was performed with 5 µL of the cloning reactions by heat shock transformation. Sequences of oligonucleotides are provided in Supplementary Table S3. Two colonies from each transformation plates were used as biological replicates in each experiment. In case of NT-CRISPR plasmids carrying multiple gRNA sequences, each gRNA expression cassette was first constructed individually on plasmids carrying a kanamycin resistance marker. A range of these plasmids is available representing the available positions in a level 2 plasmid in the framework of the Marburg Collection (Stukenberg *et al*., 2021). For this, the oligonucleotides for the respective spacers were annealed and the cloning reaction was set up as described for the NT-CRISPR plasmid above with the difference that ∼ 70 ng of plasmid DNA was used.. In a second step, those gRNA cassettes were integrated into plasmid pST_119, carrying all remaining components in a Golden Gate reaction performed as described above but with Esp3I (10,000 U/mL, NEB) instead of BsaI. The assembly of the separate gRNA expression cassettes, as well as the multi gRNA NT-CRISPR plasmid were performed using *E. coli* as a chassis, the sequence was confirmed by Sanger sequencing and the plasmids was finally introduced into *V. natriegens*.

### Preparation of tDNAs

The tDNAs used for the natural transformation were prepared by first assembling a tDNA template plasmid using Gibson Assembly (Gibson *et al*., 2009) and then use this plasmid in a PCR to generate the tDNA fragments. All tDNA templates were generated with 3 kb homologous sequences. The part entry vector of the Marburg Collection, pMC_V_01 was used as a vector for most tDNA template plasmids. In some cases, we observed a strong toxicity of the cloned sequences and used the plasmid pMC_V_11, a low-to-medium copy derivative of pMC_V_01 with p15A instead of ColE1 as the origin of replication. All primer sequences used for assembly of the tDNA template plasmids and the subsequent PCR reactions are provided in Supplementary Table S4 and S5, respectively. The template was eliminated after the PCR reactions by addition of 1 µL DpnI (10,000 U/mL, NEB) to 25 µL PCR reactions and incubation at 37 °C for at least 1 h. Lastly, a PCR cleanup was performed with the E.Z.N.A Cycle Pure Kit (Omega Bio-Tex), according to manufacturer’s instructions.

### Natural Transformation with CRISPR/Cas9 counterselection (NT-CRISPR workflow)

Natural transformation was performed largely as described previously for *V. natriegens* (Dalia *et al*., 2017) with the addition of an additional step for CRISPR/Cas9 mediated counterselection. Precultures were grown overnight (16 - 17 h) at 30 °C and 200 rpm in 5 mL LBv2 with 4 µg/mL chloramphenicol and 100 µM IPTG (Roth, CAS: 367-93-1), to induce *tfoX* expression. The natural transformation was started by adding 3.5 µL of the precultures (OD_600_ ∼ 9 - 11) to 350 µL sea salt medium (28 g/L (Sigma, S9883)) with 100 µM IPTG in a 1.5 mL reaction tube. Unless indicated otherwise, 10 ng of tDNA with 3 kb homologous flanks was added. Samples were briefly vortexed and then incubated statically at 30 °C for 5 h. In a subsequent step, 1 mL LBv2 without antibiotics was added to the cells. For CRISPR/Cas9 induction, 200 ng/mL ATc (Alfa Aeser, 13803-65-1) was added to the LBv2 medium. Tubes were mixed by inversion and incubated at 300 rpm and 30 °C for 1 h. 100 µL of appropriate dilutions (e.g. 10^−3^ for single deletions or 10^−4^ for uninduced controls) were plated on LBv2 agar plates with 2 µg/mL chloramphenicol. For CRISPR/Cas9 induction, 200 ng/mL ATc was added to the agar plates. For experiments without CRISPR/Cas9 induction, ATc was omitted from both the liquid medium and the agar plates. All experiments were performed with two biological replicates obtained from transformation of the gRNA cloning reaction in *V. natriegens* or the re-transformation of the multi-gRNA NT-CRISPR plasmid, previously assembled using *E. coli*.

### Quantification of killing efficiencies in the NT-CRISPR workflow

Samples for assessing killing efficiencies were prepared as described above for the NT-CRISPR workflow but without addition of tDNAs. For each strain, two 1.5 mL reaction tubes were run in parallel, with and without CRISPR/Cas9 counterselection. To one tube, 200 ng/mL ATc was added to the LBv2 medium and for the preparation of LBv2 agar plates and the second tube is run in parallel without any ATc. The ratio between the obtained CFU/µL from those two samples is used to calculate the killing efficiency.

### Verification of edits after NT-CRISPR

The approach used for the verification of colonies after NT-CRISPR differed between mutations types. For the quantitative characterization of NT-CRISPR (Figure 2B, 2C) with *wbfF* as the target, colonies with different morphologies were counted and used to calculate editing efficiencies. For all other targets, we tested 25 colonies from each biological replicate, resulting in 50 analyzed colonies. All deletions, apart from *wbfF*, and all integrations were verified by PCR with one primer slightly outside of the homologous flank and the other primer binding at the junction of the deletion, bridging the gap between the upstream and downstream sequence flanking the deleted region (for deletions) or binding inside the mScarlet-I sequence (for integrations). For increased throughput, PCRs were performed in 96-well plates. For this, colonies were used to inoculate 100 µL LBv2 in 96-well round bottom plates and incubated at 1000 rpm in a 96-well plate shaker for 5 - 6 h to obtain densely grown cultures. Cells were harvested by centrifugation at 3000 x g for 10 min. Medium was aspirated and cell pellets were resuspended in 100 µL H_2_0. Cell lysis was achieved by floating the 96-well plate on a water bath with 95°C for 15 min. Lastly, 96-well plates were centrifuged again at 3000 x g for 10 min and 1 µL of the supernatant was used in 12.5 µL PCR reactions with Taq polymerase (NEB) according to manufacturer’s instructions. Point mutations were confirmed phenotypically by streaking the obtained colonies on M9 minimal medium agar plates with either glucose or the alternative carbon source. Colonies growing on M9 with glucose but not with the respective secondary carbon source were considered as successfully edited. M9 agar plates were prepared (for 100 mL) by autoclaving H_2_0 (35.7 mL) together with 1.5 g agarose. Subsequently all remaining components were prewarmed to 60 °C and added to the autoclaved components as follows: 50 mL 2 x M9 salts (Na_2_HOP_4_ (17 g/L), KH_2_PO_4_ (6 g/L), NaCl (1 g/L), NH_4_Cl (2 g/L), sterile filtered), 10 mL NaCl (20 %), 4 mL carbon source (10 %), 200 µL MgSO_4_ (1 M), 100 µL (CaCl_2_). For all mutations, PCR results and phenotypic characterizations were confirmed by Sanger sequencing. Four colonies, or the maximum number of putatively correct colonies, were used for each deletion, integration or point mutation. Sequencing was performed through Microsynth Seqlab using PCR fragments and a primer binding ∼ 500 bp upstream of the respective modification. Primers used for PCR verification and sequencing are provided in Supplementary Table S6 and S7, respectively.

### Plasmid curing

After verification of the modifications, the NT-CRISPR plasmid was cured. For this, colonies yielded by the NT-CRISPR method were used to inoculate 5 mL of antibiotic free LBv2 and grown at 37 °C for 6 - 7h. To obtain single colonies, we plated 100 µL of a 10^−7^ dilution, prepared in LBV2, on antibiotic free LBv2 agar plates. After overnight incubation at 37 °C, colonies were patched on LBv2 with and without 2 µg/mL chloramphenicol to check for plasmid loss. Colonies growing on the antibiotic free agar plates but not on agar plates containing chloramphenicol were considered to be plasmid cured and glycerol stocks were prepared as described above.

### Quantification of mScarlet-I signal of *V. natriegens* reporter strains

Quantitative reporter experiments were performed largely as described before for the characterization of genetic parts (Stukenberg *et al*., 2021). Fist, material from glycerol stocks was resuspended in 50 µL LBv2 and 5 µL of the resulting suspension was used to inoculate 95 µL of LBv2 in a flat bottom 96-well plate. Cells were incubated as precultures for 5.5 - 6 h (equaling an overnight culture in similar workflows for *E. coli*) and then diluted 1:100 in fresh LBv2 to start the experiment in a Biotek Synergy H1 micro plate reader. Measurements were taken in 6 min intervals with a 3 min shaking step in double orbital mode and maximum speed occurring between measurements. The OD_600_ was measured in “normal” mode with eight measurements per data point and 100 ms delay after plate movement. mScarlet-I fluorescence was measured with a focal height of 6.5 mm, excitation and emission wavelength of 579 nm and 616 nm, respectively, and a gain of 90. Strains were used after curing of the NT-CRISPR plasmid.

### Analysis of microplate reader experiments

Sample data were first normalized by subtracting the mean of four blank wells of the background medium measurements from the sample measurements. Growth curves were computationally synchronized by aligning the first data point of each well with OD_600_ > 0.01 and the mean of the OD_600_ values of the aligned growth curves of all tested replicates and independent experiments was plotted. The relative mScarlet-I/OD_600_ values were obtained by dividing all mScarlet-I/OD_600_ values by value of the respective sample at time point 8 h, to compensate for different absolute mScarlet-I signals.

## Supporting information

Plasmid maps

## Data availability

Genetic part sequences and description of plasmid assembly are provided in Supplementary Tables S1 and S2, respectively. Sequences of all oligonucleotides used in this study are provided in Supplementary Tables S3 – S7. Plasmid maps of NT-CRISPR plasmids and plasmids carrying the separate gRNA expression cassettes for assembly of multi-gRNA NT-CRISPR plasmids are available as a Supplementary Data File. Plasmids will be made available in the future through Addgene and are available from the authors upon reasonable request. Any other relevant data are available from the corresponding author upon reasonable request.

## Acknowledgements

We thank Prof. Ankur B. Dalia for supplying plasmid pMMB-TfoX and Prof. Christopher A. Voigt for the Marionette Sensor Collection which was supplied through Addgene (Addgene Kit #1000000137). Lastly, we thank Dr. Patrick Sobetzko and Marc Teufel for intense discussions throughout the project.

## Author contributions

D.S., and A.B. conceived the design of this project. D.S. performed the majority of experiments. J.H. characterized transformation efficiency of mutant strains and A.F. constructed and tested inducible promoters used in the NT-CRISPR plasmid. D.S. analyzed the data. D.S. and A.B. wrote the manuscript. A.B. supervised the study.

## Notes

The authors declare no competing financial interest.

## References

Aiyar, S. E., Gaal, T. and Gourse, R. L. (2002) ‘rRNA promoter activity in the fast-growing bacterium Vibrio natriegens’, Journal of Bacteriology, 184(5), pp. 1349–1358. doi: 10.1128/JB.184.5.1349-1358.2002.

Aparicio, T., de Lorenzo, V. and Martínez-García, E. (2019) ‘CRISPR/Cas9-enhanced ssDNA recombineering for Pseudomonas putida’, Microbial Biotechnology, 12(5), pp. 1076–1089. doi: https://doi.org/10.1111/1751-7915.13453.

Aschenbrenner, S. et al. (2020) ‘Coupling Cas9 to artificial inhibitory domains enhances CRISPR-Cas9 target specificity’, Science Advances, 6(6), p. eaay0187. doi: 10.1126/sciadv.aay0187.

Baba, T. et al. (2006) ‘Construction of Escherichia coli K-12 in-frame, single-gene knockout mutants: the Keio collection’, Molecular systems biology. 2006/02/21, 2, pp. 2006.0008-2006.0008. doi: 10.1038/msb4100050.

Bik, E. M. et al. (1996) ‘Genetic organization and functional analysis of the otn DNA essential for cell-wall polysaccharide synthesis in Vibrio cholerae O139’, Molecular Microbiology, 20(4), pp. 799–811. doi: https://doi.org/10.1111/j.1365-2958.1996.tb02518.x.

Bindels, D. S. et al. (2017) ‘mScarlet: a bright monomeric red fluorescent protein for cellular imaging.’, Nature methods, 14(1), pp. 53–56. doi: 10.1038/nmeth.4074.

Blokesch, M. and Schoolnik, G. K. (2008) ‘The Extracellular Nuclease Dns and Its Role in Natural Transformation of Vibrio cholerae’, Journal of Bacteriology, 190(21), pp. 7232LP–7240. doi: 10.1128/JB.00959-08.

Blomqvist, T., Steinmoen, H. and Håvarstein, L. S. (2006) ‘Natural genetic transformation: A novel tool for efficient genetic engineering of the dairy bacterium Streptococcus thermophilus.’, Applied and environmental microbiology, 72(10), pp. 6751–6756. doi: 10.1128/AEM.01156-06.

Bubeck, F. et al. (2018) ‘Engineered anti-CRISPR proteins for optogenetic control of CRISPR–Cas9’, Nature Methods, 15(11), pp. 924–927. doi: 10.1038/s41592-018-0178-9.

Bursztyn, H. et al. (1975) ‘Transfectability of rough strains of Salmonella typhimurium.’, Journal of bacteriology, 124(3), pp. 1630–1634. doi: 10.1128/JB.124.3.1630-1634.1975.

Cavanagh, A. T., Chandrangsu, P. and Wassarman, K. M. (2010) ‘6S RNA regulation of relA alters ppGpp levels in early stationary phase’ Microbiology (Reading, England). 2010/09/09, 156(Pt 12), pp. 3791–3800. doi: 10.1099/mic.0.043992-0.

Cui, L. and Bikard, D. (2016) ‘Consequences of Cas9 cleavage in the chromosome of Escherichia coli’, Nucleic acids research. 2016/04/08, 44(9), pp. 4243–4251. doi: 10.1093/nar/gkw223.

Dalia, A. B., McDonough, E. and Camilli, A. (2014) ‘Multiplex genome editing by natural transformation’, Proceedings of the National Academy of Sciences, 111(24), pp. 8937LP–8942. doi: 10.1073/pnas.1406478111.

Dalia Triana, N et al. (2017) ‘Enhancing multiplex genome editing by natural transformation (MuGENT) via inactivation of ssDNA exonucleases’, Nucleic acids research, 45(12), pp. 7527–7537. doi: 10.1093/nar/gkx496.

Dalia Triana, N. et al. (2017) ‘Multiplex Genome Editing by Natural Transformation (MuGENT) for Synthetic Biology in Vibrio natriegens’ ACS Synthetic Biology, 6(9), pp. 1650–1655. doi: 10.1021/acssynbio.7b00116.

Detloff, P., Sieber, J. and Petes, T. D. (1991) ‘Repair of specific base pair mismatches formed during meiotic recombination in the yeast Saccharomyces cerevisiae.’, Molecular and Cellular Biology, 11(2), pp. 737LP–745. doi: 10.1128/MCB.11.2.737.

Van Die, I. M. et al. (1984) ‘Transformability of galE variants derived from uropathogenic Escherichia coli strains’, Journal of Bacteriology, 158(2), pp. 760–761. doi: 10.1128/jb.158.2.760-761.1984.

Eagon, R. G. (1961) ‘Generation Time of Less Than 10 Minutes’, pp. 1961–1962.

Gibson, D. G. et al. (2009) ‘Enzymatic assembly of DNA molecules up to several hundred kilobases’, Nature Methods, 6(5), pp. 343–345. doi: 10.1038/nmeth.1318.

Hoff, J. et al. (2020) ‘Vibrio natriegens: an ultrafast-growing marine bacterium as emerging synthetic biology chassis’, Environmental Microbiology, 22(10), pp. 4394–4408. doi: 10.1111/1462-2920.15128.

Hoffart, E. et al. (2017) ‘High substrate uptake rates empower Vibrio natriegens as production host for industrial biotechnology’, Applied and Environmental Microbiology, 83(22), pp. 1–10. doi: 10.1128/AEM.01614-17.

Hoffmann, M. D. et al. (2019) ‘Cell-specific CRISPR-Cas9 activation by microRNA-dependent expression of anti-CRISPR proteins.’, Nucleic acids research, 47(13), p. e75. doi: 10.1093/nar/gkz271.

Hu, J. H. et al. (2018) ‘Evolved Cas9 variants with broad PAM compatibility and high DNA specificity.’, Nature, 556(7699), pp. 57–63. doi: 10.1038/nature26155.

Isaacs, F. J. et al. (2011) ‘Precise Manipulation of Chromosomes in Vivo Enables Genome-Wide Codon Replacement’, Science, 333(6040), pp. 348LP–353. doi: 10.1126/science.1205822.

Jaishankar, J. and Srivastava, P. (2017) ‘Molecular Basis of Stationary Phase Survival and Applications ‘, Frontiers in Microbiology, p. 2000. Available at: https://www.frontiersin.org/article/10.3389/fmicb.2017.02000.

Jiang, W. et al. (2013) ‘RNA-guided editing of bacterial genomes using CRISPR-Cas systems’, Nature Biotechnology, 31(3), pp. 233–239. doi: 10.1038/nbt.2508.

Jiang, Y. et al. (2015) ‘Multigene Editing in the Escherichia coli Genome via the CRISPR-Cas9 System’, Applied and Environmental Microbiology. edited by R.M. Kelly, 81(7), pp. 2506LP–2514. doi: 10.1128/AEM.04023-14.

Johnsborg, O., Eldholm, V. and Håvarstein, L. S. (2007) ‘Natural genetic transformation: prevalence, mechanisms and function’, Research in Microbiology, 158(10), pp. 767–778. doi: https://doi.org/10.1016/j.resmic.2007.09.004.

Kato, J. and Hashimoto, M. (2007) ‘Construction of consecutive deletions of the Escherichia coli chromosome’, Molecular systems biology. 2007/08/14, 3, p. 132. doi: 10.1038/msb4100174.

Kim, I. et al. (2018) ‘Solution structure and dynamics of anti-CRISPR AcrIIA4, the Cas9 inhibitor’, Scientific Reports, 8(1), p. 3883. doi: 10.1038/s41598-018-22177-0.

Kleinstiver, B. P. et al. (2015) ‘Engineered CRISPR-Cas9 nucleases with altered PAM specificities’, Nature, 523(7561), pp. 481–485. doi: 10.1038/nature14592.

Lee, H. H. et al. (2016) ‘Vibrio natriegens, a new genomic powerhouse’, bioRxiv, p. 58487. doi:10.1101/058487.

Lee, H. H. et al. (2017) ‘Recombineering in Vibrio natriegens’, bioRxiv, p. 130088. doi: 10.1101/130088.

Lee, H. H. et al. (2019) ‘Functional genomics of the rapidly replicating bacterium Vibrio natriegens by CRISPRi’, Nature Microbiology, 4(7), pp. 1105–1113. doi: 10.1038/s41564-019-0423-8.

Lee, J. H. et al. (2009) ‘Metabolic engineering of a reduced-genome strain of Escherichia coli for L-threonine production’, Microbial Cell Factories, 8(1), p. 2. doi: 10.1186/1475-2859-8-2.

Lieber, M. R. (2010) ‘The mechanism of double-strand DNA break repair by the nonhomologous DNA end-joining pathway’, Annual review of biochemistry, 79, pp. 181–211. doi: 10.1146/annurev.biochem.052308.093131.

Maida, I. et al. (2013) ‘Draft Genome Sequence of the Fast-Growing Bacterium Vibrio natriegens Strain DSMZ 759.’, Genome announcements, 1(4). doi: 10.1128/genomeA.00648-13.

Meier, P. and Wackernagel, W. (2005) ‘Impact of mutS inactivation on foreign DNA acquisition by natural transformation in Pseudomonas stutzeri’, Journal of bacteriology, 187(1), pp. 143–154. doi: 10.1128/JB.187.1.143-154.2005.

Meyer, A. J. et al. (2019) ‘Escherichia coli “Marionette” strains with 12 highly optimized small-molecule sensors.’, Nature chemical biology, 15(2), pp. 196–204. doi: 10.1038/s41589-018-0168-3.

Nikolay, R. et al. (2014) ‘Validation of a fluorescence-based screening concept to identify ribosome assembly defects in Escherichia coli’ Nucleic Acids Research, 42(12), pp. e100–e100. doi: 10.1093/nar/gku381.

Payne, W. J. (1958) ‘Studies on bacterial utilization of uronic acids. III. Induction of oxidative enzymes in a marine isolate.’, Journal of bacteriology, 76(3), pp. 301–307. doi: 10.1128/JB.76.3.301-307.1958.

Penewit, K. et al. (2018) ‘Efficient and Scalable Precision Genome Editing in Staphylococcus aureus through Conditional Recombineering and CRISPR/Cas9-Mediated Counterselection’, mBio, 9(1), pp. e00067–18. doi: 10.1128/mBio.00067-18.

Pfeifer, E. et al. (2019) ‘Generation of a Prophage-Free Variant of the Fast-Growing Bacterium Vibrio natriegens.’, Applied and environmental microbiology, 85(17). doi: 10.1128/AEM.00853-19.

Pósfai, G. et al. (2006) ‘Emergent Properties of Reduced-Genome Escherichia coli’, Science, 312(5776), pp. 1044LP–1046. doi: 10.1126/science.1126439.

Reisch, C. R. and Prather, K. L. J. (2015) ‘The no-SCAR (Scarless Cas9 Assisted Recombineering) system for genome editing in Escherichia coli’, Scientific Reports, 5(1), p. 15096. doi: 10.1038/srep15096.

Seitz, P. and Blokesch, M. (2013) ‘Cues and regulatory pathways involved in natural competence and transformation in pathogenic and environmental Gram-negative bacteria’, FEMS Microbiology Reviews, 37(3), pp. 336–363. doi: 10.1111/j.1574-6976.2012.00353.x.

Selas Castiñeiras, T. et al. (2018) ‘E. coli strain engineering for the production of advanced biopharmaceutical products’, FEMS Microbiology Letters, 365(15). doi: 10.1093/femsle/fny162.

Shin, J. et al. (2017) ‘Disabling Cas9 by an anti-CRISPR DNA mimic.’, Science advances, 3(7), p. e1701620. doi: 10.1126/sciadv.1701620.

Shuman, S. and Glickman, M. S. (2007) ‘Bacterial DNA repair by non-homologous end joining’, Nature Reviews Microbiology, 5(11), pp. 852–861. doi: 10.1038/nrmicro1768.

Stukenberg, D. et al. (2021) ‘The Marburg Collection: A Golden Gate DNA Assembly Framework for Synthetic Biology Applications in Vibrio natriegens.’, ACS synthetic biology. doi: 10.1021/acssynbio.1c00126.

Su, S. S. et al. (1988) ‘Mispair specificity of methyl-directed DNA mismatch correction in vitro.’, Journal of Biological Chemistry, 263(14), pp. 6829–6835. doi: https://doi.org/10.1016/S0021-9258(18)68718-6.

Visick, K. L. et al. (2018) ‘Tools for Rapid Genetic Engineering of Vibrio fischeri.’, Applied and environmental microbiology, 84(14). doi: 10.1128/AEM.00850-18.

Vojcic, L. et al. (2012) ‘An efficient transformation method for Bacillus subtilis DB104’, Applied Microbiology and Biotechnology, 94(2), pp. 487–493. doi: 10.1007/s00253-012-3987-2.

Walton, R. T. et al. (2020) ‘Unconstrained genome targeting with near-PAMless engineered CRISPR-Cas9 variants’, Science, 368(6488), pp. 290LP–296. doi: 10.1126/science.aba8853.

Wang, H. H. et al. (2009) ‘Programming cells by multiplex genome engineering and accelerated evolution’, Nature. 2009/07/26, 460(7257), pp. 894–898. doi: 10.1038/nature08187.

Wang, Z. et al. (2013) ‘Draft Genome Sequence of the Fast-Growing Marine Bacterium Vibrio natriegens Strain ATCC 14048’, Genome announcements, 1(4), pp. e00589–13. doi: 10.1128/genomeA.00589-13.

Wasels, F. et al. (2020) ‘A CRISPR/Anti-CRISPR Genome Editing Approach Underlines the Synergy of Butanol Dehydrogenases in Clostridium acetobutylicum DSM 792’, Applied and environmental microbiology, 86(13), pp. e00408–20. doi: 10.1128/AEM.00408-20.

Weinstock, M. T. et al. (2016) ‘Vibrio natriegens as a fast-growing host for molecular biology’, Nature Methods, 13(10), pp. 849–851. doi: 10.1038/nmeth.3970.

Yang, H. and Patel, D. J. (2017) ‘Inhibition Mechanism of an Anti-CRISPR Suppressor AcrIIA4 Targeting SpyCas9.’, Molecular cell, 67(1), pp. 117-127.e5. doi: 10.1016/j.molcel.2017.05.024.

