## Supplementary Figures and Tables for "NT-CRISPR: Combining natural transformation and CRISPR/Cas9 counterselection for markerless and scarless genome editing in *Vibrio natriegens*"

### Contents

Supplementary Figure S1 - Stoichiometry of Cas9 and AcrIIA4 is essential for inducible cell killing.

Supplementary Figure S2 - Underlying CFU/µL for Figure 2B and 2C

Supplementary Figure S3 - Sequencing results of Δ*vnp1* strains

Supplementary Figure S4 - Transformation efficiency of deletion strains

Supplementary Figure S5 - Assembly of NT-CRISPR plasmid with multiple gRNAs

Supplementary Figure S6 - Killing assay with strains carrying plasmids with three gRNAs

Supplementary Figure S7 - mScarlet-I/$OD_{600}$ signal of strains shown in Figure 4D

Supplementary Table S1 - New genetic parts generated in this study

Supplementary Table S2 - Assembly of plasmids used in this study

Supplementary Table S3 - Oligonucleotides used to assemble gRNA sequences

Supplementary Table S4 - Oligonucleotides used for the construction of tDNA template plasmids

Supplementary Table S5 - Oligonucleotides to generate tDNA fragments from tDNA template plasmids

Supplementary Table S6 - Oligonucleotides used for PCR screening of deletions and integrations

Supplementary Table S7 - Oligonucleotides used for PCR to amplify target region for sequencing

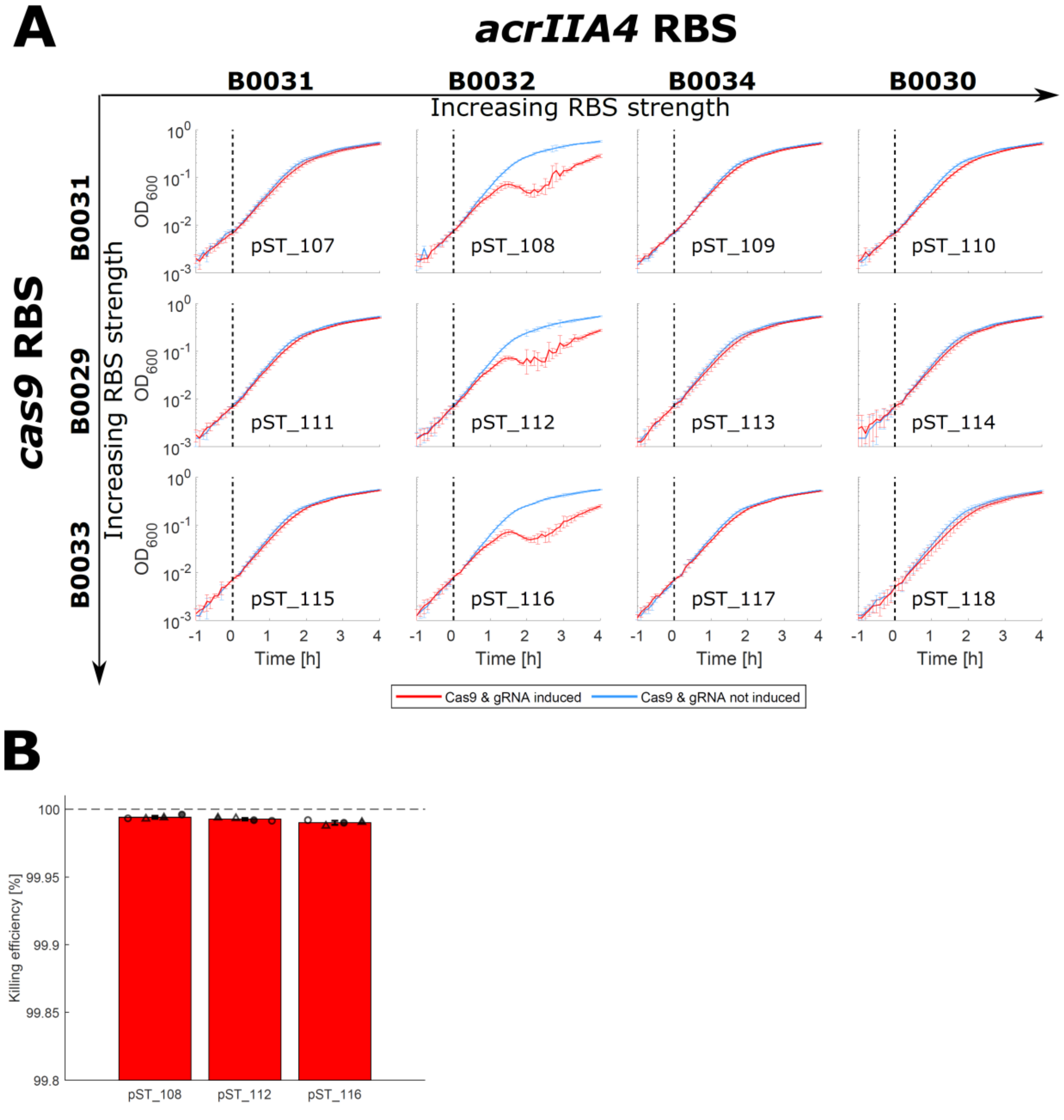


*Supplementary Figure S1: Stoichiometry of Cas9 and AcrIIA4 is essential for inducible cell killing. (A) Inducible cell killing in liquid culture, measured in a microplate reader. Constructs carry combinations of different RBS for cas9 and acrIIA4, provided on top and left side and a gRNA targeting wbfF. Strength of RBS, based on characterization experiments with fluorescent reporter genes* (Stukenberg *et al.*, 2021)*, increases from left to right and top to bottom. Data represent mean of two biological replicates and two independent experiments. Error bars show the standard deviation from the mean. Samples were induced with 200 ng/µL ATc (red) or remained uninduced (blue) at time point 0 (dashed lines). (B) Results of killing assay in the NT-CRISPR workflow. Killing efficiency is calculated as follows:* $Killing\ efficiency\ [\%] = 1 - \frac{CFU/\mu L\ with\ counterselection}{CFU/\mu L\ without\ counterselection}$ ** 100. Data represent the mean of two biological replicates (circle or triangle) and two independent experiments (filled or open symbols). Error bars indicate standard deviation of the mean. Dashed line indicates the highest possible value.*

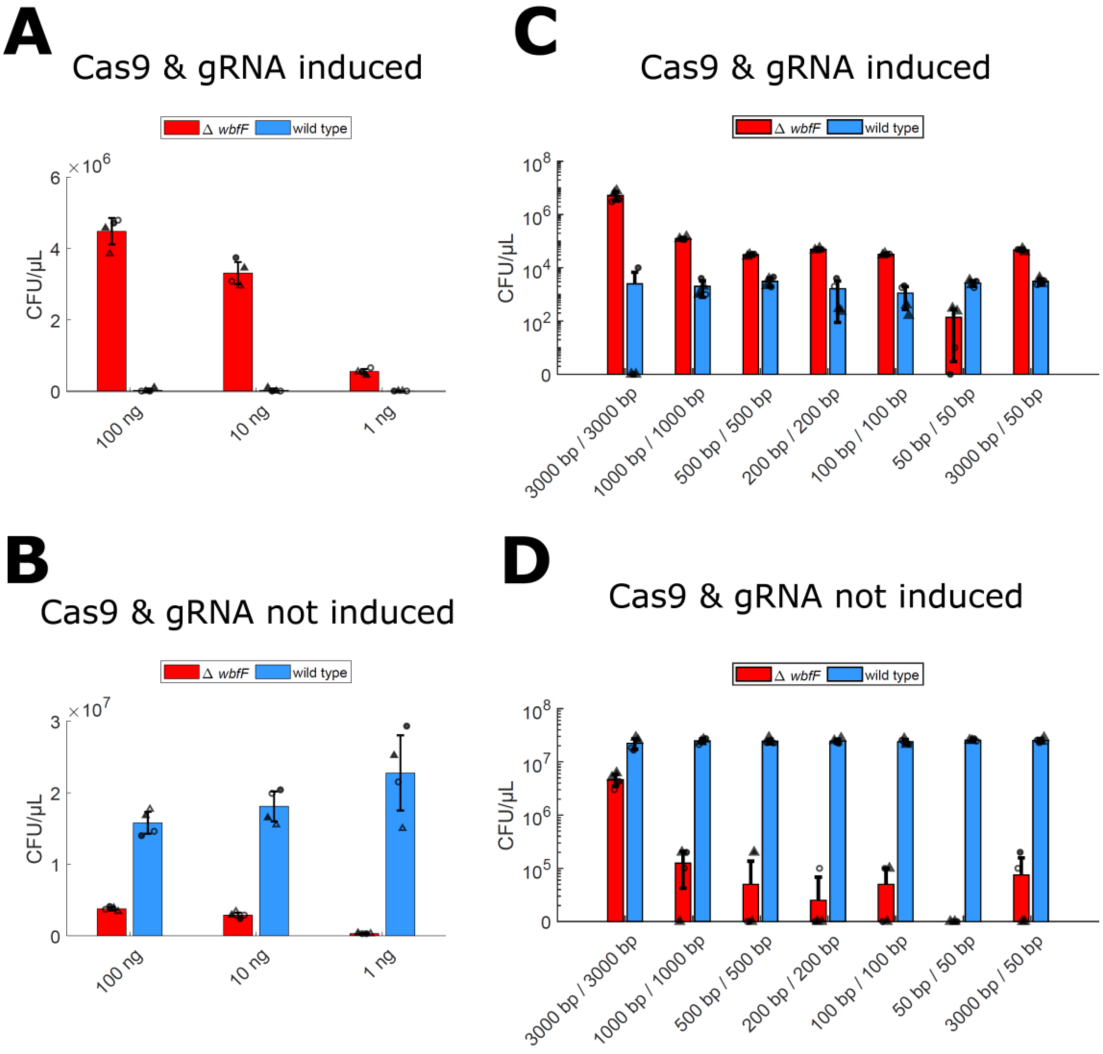


*Supplementary Figure S2: Underlying CFU/µL for Figure 2B and 2C. Red and blue bars show CFU/µL for colonies with the transparent ΔwbfF morphology (red bars) or wild type morphology (blue bars). Data represent the mean of two biological replicates (circle or triangle) and two independent experiments (filled or open symbols). Error bars indicate standard deviation of the mean. (A,B) CFU/µL of experiment with different amounts of tDNA with 3000 bp homologous flanks upstream and downstream of the target sequence. (C,D) CFU/µL of experiment with 100 ng of tDNA with different length of homologous flanks. (A,C) Results with CRISPR-based counterselection. (B,D) Results without CRISPR-based counterselection.*

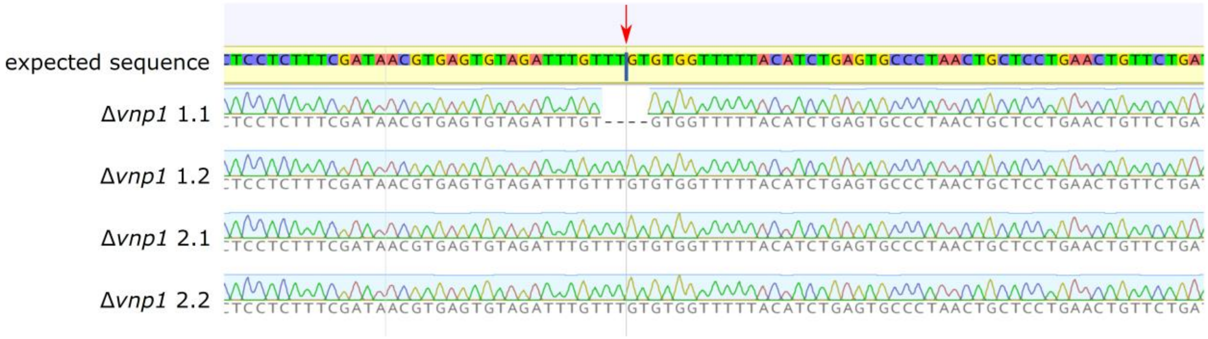


*Supplementary Figure S3: Sequencing results of Δvnp1 strains. The expected sequence for a Δvnp1 deletion is shown on top. The red arrow indicates the junction between upstream and downstream sequence surrounding the deleted fragments. Δvnp1 clone 1.1 shows a deviation from the expected sequence with four missing bases. Sequence alignment was created with Geneious Prime 2021.2.1.*

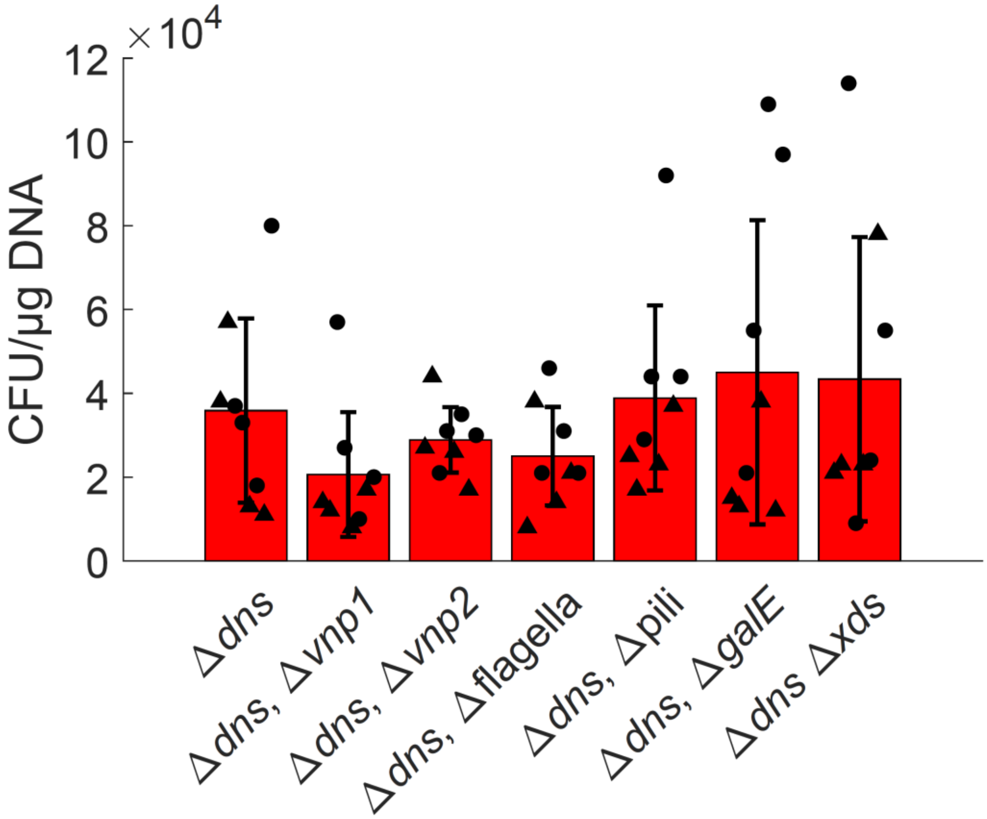


*Supplementary Figure S4: Transformation efficiency of deletion strains. Parental strain Δdns and derivatives with additional deletions were tested for transformation efficiency. CFU/µg DNA was determined from four replicates in two independent experiments (circles and triangles). The plasmid pMC0_8_19* (Stukenberg *et al.*, 2021)*, was used for this experiment.*

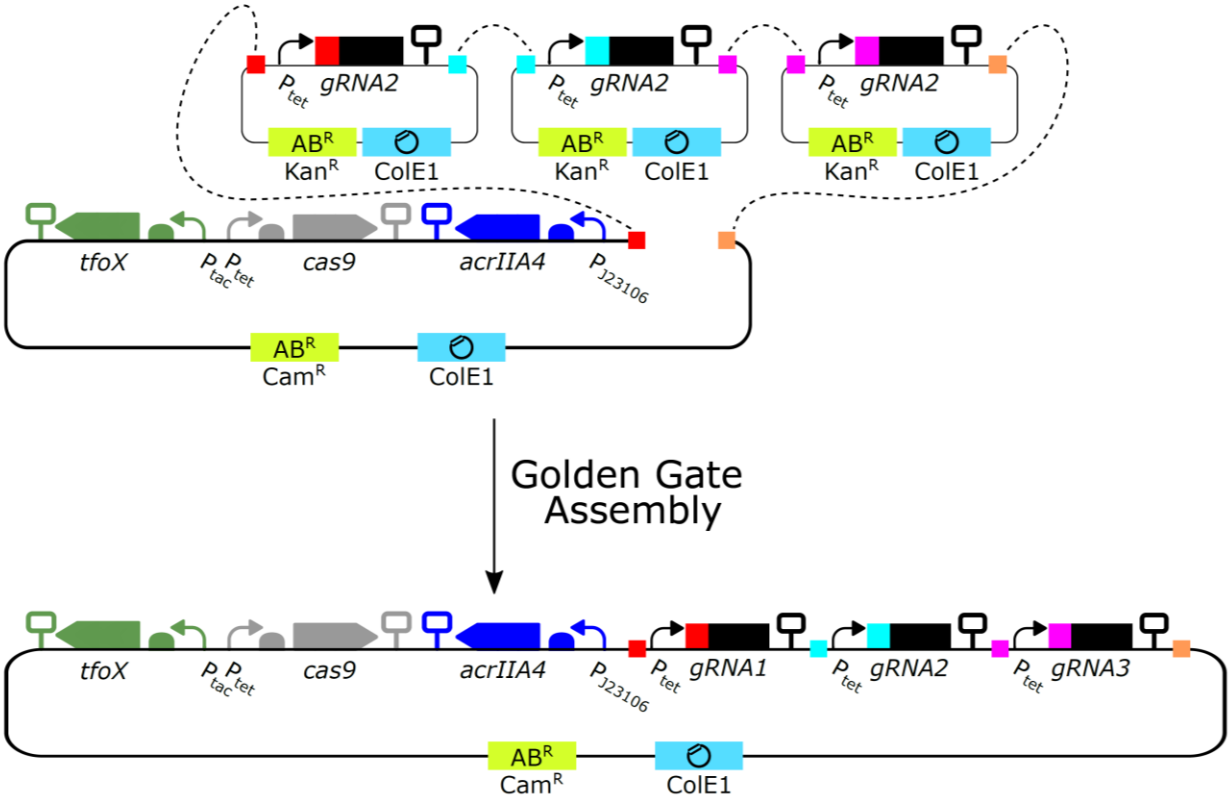


*Supplementary Figure S5: Assembly of NT-CRISPR plasmid with multiple gRNAs: First individual gRNA expression cassettes are created in plasmids carrying a kanamycin resistance marker (*Kan$^R$*) and then subsequently integrated into a plasmid carrying all remaining components by Golden Gate Assembly with Esp3I as the type IIs restriction enzyme. Colored squares indicate matching fusion sites for Golden Gate Assembly.*

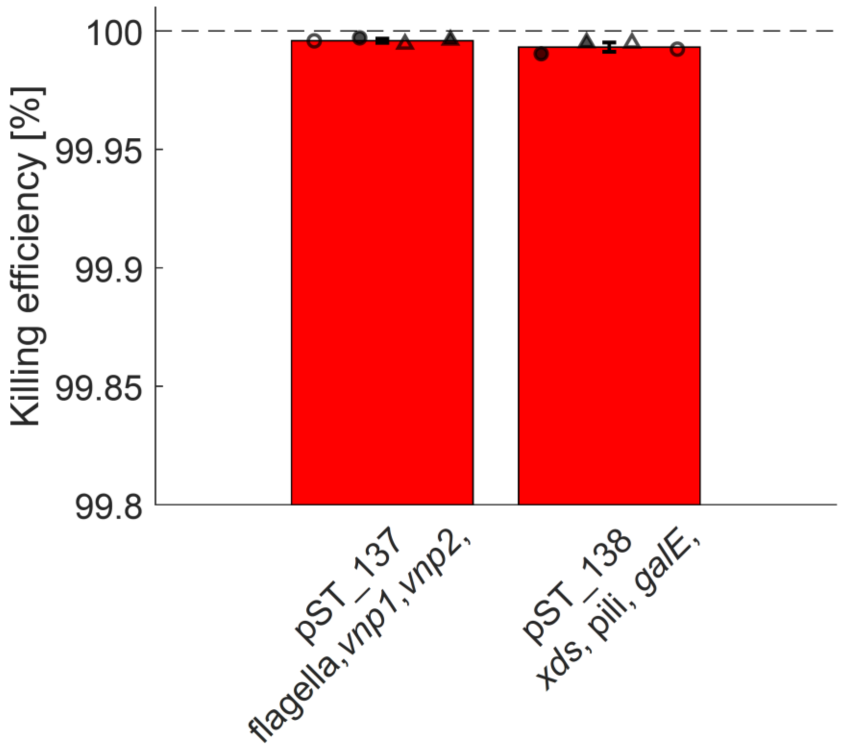


*Supplementary Figure S6: Killing assay with strains carrying plasmids with three gRNAs. Killing efficiency is calculated as follows* $Killing\ efficiency\ [\%] = 1 - \frac{CFU/\mu L\ with\ counterselection}{CFU/\mu L\ without\ counterselection}$ ** 100. Data represent the mean of two biological replicates (circle or triangle) and two independent experiments (filled or open symbols). Error bars indicate standard deviation of the mean. Dashed line indicates the highest possible value.*

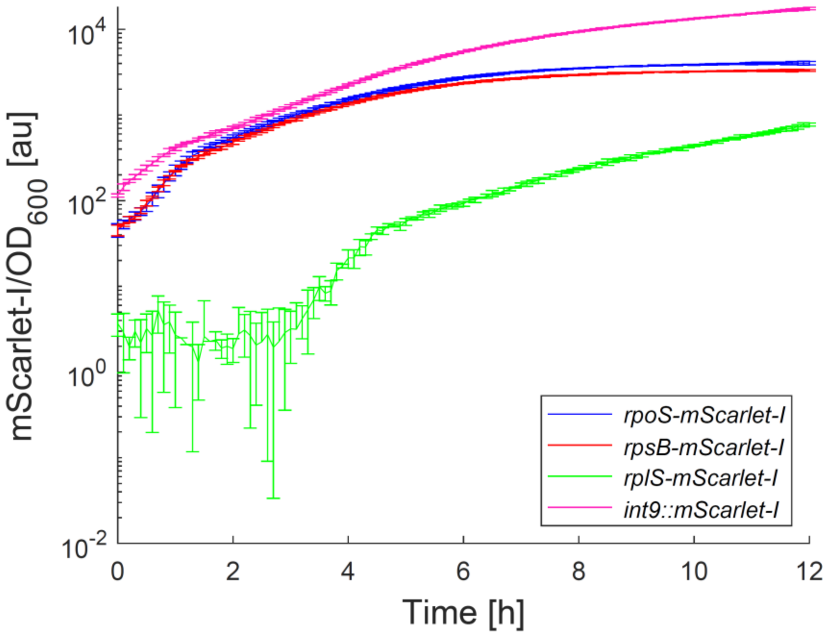


*Supplementary Figure S6: mScarlet-I/OD$_{600}$ signal of strains shown in Figure 4D. Data represent the mean of four biological replicates. Error bars indicate standard deviation from the mean.*

### Supplementary Tables

*Supplementary Table S1: New genetic parts generated in this study. Parts were assembled as described for the Marburg Collection* (Stukenberg *et al.*, 2021). *First and last four bases represent fusion sites for the subsequent assembly.*

| Part | Sequence |
|---|---|
| pMC0_2_42_Ptet* | GGAGCTCAGATAAAATATTTGCTCATGAGCCCGAAGTGGCGAGCCCGATCTTCCCCATCGGTGATGTCGGCGATATAGGCGCCA<br>GCAACCGCACCTGTGGCGCCGGTGATGCCGGCCACGATGCGTCCGGCGTAGAGGATCTGCTCATGTTTGACAGCTTATCATCGA<br>TGCATAATGTGCCTGTCAAATGGACGAAGCAGGGATTCTGCAAACCCTATGCTACTCCCTCGAGCCGTCAATTCTGATTCGTTACC<br>AATTAGGATCCTTATCAGGACCCACTTTCACATTTAAGTTGTTTTTCTAATCCGCATATGATCAATTCAAGGCCGAATAAGAAGGC<br>TGGCTCTGCACCTTGGTGATCAAATAATTCGATAGCTTGTCGTAATAATGGCGGCATACTATCAGTAGTAGGTGTTTCCCTTTCTT<br>CTTTAGCGACTTGATGCTCTTGATCTTCCAATACGCAACCTAAAGTAAAATGCCCCACAGCGCTGAGTGCATATAATGCATTCTCT<br>AGTGAAAAACCTTGTTGGCATAAAAAGGCTAATTGATTTTCGAGAGTTTCATACTGTTTTTCTGTAGGCCGTGTACCTAAATGTAC<br>TTTTGCTCCATCGCGATGACTTAGTAAAGCACATCTAAAACTTTTAGCGTTATTACGTAAAAAATCTTGCCAGCTTTCCCCTTCTAA<br>AGGGCAAAAGTGAGTATGGTGCCTATCTAACATCTCAATGGCTAAGGCGTCGAGCAAAGCCCGCTTATTTTTTACATGCCAATAC<br>AATGTAGGCTGCTCTACACCTAGCTTCTGGGCGAGTTTACGGGTTGTTAAACCTTCGATTCCGACCTCATTAAGCAGCTCTAATGC<br>GCTGTTAATCACTTTACTTTTATCTAATCTGGACATATTCACCACCCTGAATTGACTCTCTTCCGGGCGCTATCATGCCATACCGCG<br>AAAGGTTTTGCGCCATTCGGCTGTTTTCAGCAGGACGCACTGACCTCCCTATCAGTGATAGAGATTGACATCCCTATCAGTGATA<br>GAGATACTGAGCACTACT |
| pMC0_2_43_Ptac | GGAGCTCAGATAAAATATTTGCTCATGAGCCCGAAGTGGCGAGCCCGATCTTCCCCATCGGTGATGTCGGCGATATAGGCGCCA<br>GCAACCGCACCTGTGGCGCCGGTGATGCCGGCCACGATGCGTCCGGCGTAGAGGATCTGCTCATGTTTGACAGCTTATCATCGA<br>TGCATAATGTGCCTGTCAAATGGACGAAGCAGGGATTCTGCAAACCCTATGCTACTCCCTCGAGCCGTCAATTGTCTGATTCGTTA<br>CCAATTAGGATCCTTATCACTGCCCGCTTTCCAGTCGGGAAACCTGTCGTGCCAGCTGCATTAATGAATCGGCCAACGCGCGGGG<br>AGAGGCGGTTTGCGTATTGGGCGCCAGGGTGGTTTTTCTTTTCACCAGTGAGACTGGCAACAGCTGATTGCCCTTCACCGCCTGG<br>CCCTGAGAGAGTTGCAGCAAGCGGTCCACGCTGGTTTGCCCCAGCAGGCGAAAATCCTGTTTGATGGTGGTTAACGGCGGGATA<br>TAACATGAGCTATCTTCGGTATCGTCGTATCCCACTACCGAGATATCCGCACCAACGCGCAGCCCGGACTCGGTAATGGCGCGCA<br>TTGCGCCCAGCGCCATCTGATCGTTGGCAACCAGCATCGCAGTGGGAACGATGCCCTCATTCAGCATTTGCATGGTTTGTTGAAA<br>ACCGGACATGGCACTCCAGTCGCCTTCCCGTTCCGCTATCGGCTGAATTTGATTGCGAGTGAGATATTTATGCCAGCCAGCCAGA<br>CGCAGACGCGCCGAGACAGAACTTAATGGGCCCGCTAACAGCGCGATTTGCTGGTGACCCAATGCGACCAGATGCTCCACGCCC<br>AGTCGCGTACCGTCCTCATGGGAGTAAATAATACTGTTGATGGGTGTCTGGTCAGAGACATCAAGAAATAACGCCGGAACATTA<br>GTGCAGGCAGCTTCCACAGCAATGGCATCCTGGTCATCCAGCGGATAGTTAATGATCAGCCCACTGACGCGTTGCGCGAGAAGA<br>TTGTGCACCGCCGCTTTACAGGCTTCGACGCCGCTTCGTTCTACCATCGACACCACCACGCTGGCACCCAGTTGATCGGCGCGAG<br>ATTTAATCGCCGCGACAATTTGCGACGGCGCGTGCAGGGCCAGACTGGAGGTGGCAACGCCAATCAGCAACGACTGTTTGCCCG<br>CCAGTTGTTGTGCCACGCGGTTGGGAATGTAATTCAGCTCCACCATCGCCGCTTCCACTTTTTCCCGCGTTTTCGCAGAAACGTGG<br>CTGGCCTGGTTCACCACGCGGGAAACGGTCATATAAGAGACACCGGCATACTCTGCGACATCGTATAACGTTACTGGTTTCATAT<br>TCACCACCCTGAATTGACTCTCTTCCGGGCGCTATCATGCCATACCGCGAAAGGTTTTGCGCCATTCGGCTGTGTTGACAATTAAT<br>CATCGGCTCGTATAATGTGTGGAATTGTGAGCGCTCACAATTTACT |
| pMC0_4_25_CDSCas9 | AATGGATAAGAAATACTCAATAGGCTTAGATATCGGCACAAATAGCGTCGGATGGGCGGTGATCACTGATGAATATAAGGTTCC<br>GTCTAAAAAGTTCAAGGTTCTGGGAAATACAGACCGCCACAGTATCAAAAAAAATCTTATAGGGGCTCTTTTATTTGACAGTGGA<br>GAGACAGCGGAAGCGACTCGTCTGAAACGGACAGCTCGTAGAAGGTATACACGTCGGAAGAATCGTATTTGTTATCTACAGGAG<br>ATTTTTTCAAATGAGATGGCGAAAGTAGATGATAGTTTCTTTCATCGACTTGAAGAGTCTTTTTTGGTGGAAGAAGACAAGAAGC<br>ATGAACGTCATCCTATTTTTGGAAATATAGTAGATGAAGTTGCTTATCATGAGAAATATCCAACTATCTATCATCTGCGAAAAAAA<br>TTGGTAGATTCTACTGATAAAGCGGATTTGCGCTTAATCTATTTGGCCTTAGCGCATATGATTAAGTTTCGTGGTCATTTTTTGATT<br>GAGGGAGATTTAAATCCTGATAATAGTGATGTGGACAAACTATTTATCCAGTTGGTACAAACCTACAATCAATTATTTGAAGAAA<br>ACCCTATTAACGCAAGTGGAGTAGATGCTAAAGCGATTCTTTCTGCACGATTGAGTAAATCAAGACGATTAGAAAATCTCATTGC<br>TCAGCTCCCCGGTGAGAAGAAAAATGGCTTATTTGGGAATCTCATTGCTTTGTCATTGGGTTTGACCCCTAATTTTAAATCAAATT<br>TTGATTTGGCAGAAGATGCTAAATTACAGCTTTCAAAAGATACTTACGATGATGATTTAGATAATTTATTGGCGCAAATTGGAGAT<br>CAATATGCTGATTTGTTTTTGGCAGCTAAGAATTTATCAGATGCTATTTTACTTTCAGATATCCTAAGAGTAAATACTGAAATAACT<br>AAGGCTCCCCTATCAGCTTCAATGATTAAACGCTACGATGAACATCATCAAGACTTGACTCTTTTAAAAGCTTTAGTTCGACAACA<br>ACTTCCAGAAAAGTATAAAGAAATCTTTTTTGATCAATCAAAAAACGGATATGCAGGTTATATTGATGGGGGAGCTAGCCAAGAA<br>GAATTTTATAAATTTATCAAACCAATTTTAGAAAAAATGGATGGTACTGAGGAATTATTGGTGAAACTAAATCGTGAAGATTTGCT<br>GCGCAAGCAACGGACCTTTGACAACGGCTCTATTCCCCATCAAATTCACTTGGGTGAGCTGCATGCTATTTTGAGAAGACAAGAA<br>GACTTTTATCCATTTTTAAAAGACAATCGTGAGAAGATTGAAAAAATCTTGACTTTTCGTATTCCTTATTATGTTGGTCCATTGGCG<br>CGTGGCAATAGTCGTTTTGCATGGATGACTCGGAAGTCTGAAGAAACAATTACCCCATGGAATTTTGAAGAAGTTGTCGATAAAG<br>GTGCTTCAGCTCAATCATTTATTGAACGCATGACAAACTTTGATAAAAATCTTCCAAATGAAAAAGTACTACCAAAACATAGTTTG<br>CTTTATGAGTATTTTACGGTTTATAACGAATTGACAAAGGTCAAATATGTTACTGAAGGAATGCGAAAACCAGCATTTCTTTCAGG<br>TGAACAGAAGAAAGCCATTGTTGATTTACTCTTCAAAACAAATCGAAAAGTAACCGTTAAGCAATTAAAAGAAGATTATTTCAAA<br>AAAATAGAATGTTTTGATAGTGTTGAAATTTCAGGAGTTGAAGATAGATTTAATGCTTCATTAGGTACCTACCATGATTTGCTAAA<br>AATTATTAAAGATAAAGATTTTTTGGATAATGAAGAAAATGAAGATATCTTAGAGGATATTGTTTTAACATTGACCTTATTTGAAG<br>ATAGGGAGATGATTGAGGAAAGACTTAAAACATATGCTCACCTCTTTGATGATAAGGTGATGAAACAGCTTAAACGTCGCCGTTA<br>TACTGGTTGGGGACGTTTGTCTCGAAAATTGATTAATGGTATTAGGGATAAGCAATCTGGCAAAACAATATTAGATTTTTTGAAA<br>TCAGATGGTTTTGCCAATCGCAATTTTATGCAGCTGATCCATGATGATAGTTTGACATTTAAAGAAGACATTCAAAAAGCACAAGT<br>GTCTGGACAAGGCGATAGTTTACATGAACATATTGCAAATTTAGCTGGTAGCCCTGCTATTAAAAAAGGTATTTTACAGACTGTA<br>AAAGTTGTTGATGAATTGGTCAAAGTAATGGGGCGGCATAAGCCAGAAAATATCGTTATTGAAATGGCACGTGAAAATCAGACA<br>ACTCAAAAGGGCCAGAAAAATTCGCGAGAGCGTATGAAACGAATCGAAGAAGGTATCAAAGAATTAGGAAGTCAGATTCTTAA |

| | |
|---|---|
| | AGAGCATCCTGTTGAAAATACTCAATTGCAAAATGAAAAGCTCTATCTCTATTATCTCCAAAATGGAAGAGACATGTATGTGGAC<br>CAAGAATTAGATATTAATCGTTTAAGTGATTATGATGTCGATCACATTGTTCCACAAAGTTTCCTTAAAGACGATTCAATAGACAA<br>TAAGGTCTTAACGCGTTCTGATAAAAATCGTGGTAAATCGGATAACGTTCCAAGTGAAGAAGTAGTCAAAAAGATGAAAAACTA<br>TTGGAGACAACTTCTAAACGCCAAGTTAATCACTCAACGTAAGTTTGATAATTTAACGAAAGCTGAACGTGGAGGTTTGAGTGAA<br>CTTGATAAAGCTGGTTTTATCAAACGCCAATTGGTTGAAACTCGCCAAATCACTAAGCATGTGGCACAAATTTTGGATAGTCGCAT<br>GAATACTAAATACGATGAAAATGATAAACTTATTCGAGAGGTTAAAGTGATTACCTTAAAATCTAAATTAGTTTCTGACTTCCGAA<br>AAGATTTCCAATTCTATAAAGTACGTGAGATTAACAATTACCATCATGCCCATGATGCGTATCTAAATGCCGTCGTTGGAACTGCT<br>TTGATTAAGAAATATCCAAAACTTGAATCGGAGTTTGTCTATGGTGATTATAAAGTTTATGATGTTCGTAAAATGATTGCTAAGTC<br>TGAGCAAGAAATAGGCAAAGCAACCGCAAAATATTTCTTTTACTCTAATATCATGAACTTCTTCAAAACAGAAATTACACTTGCAA<br>ATGGAGAGATTCGCAAACGCCCTCTAATCGAAACTAATGGGGAAACTGGAGAAATTGTCTGGGATAAAGGGCGAGATTTTGCCA<br>CAGTGCGCAAAGTATTGTCCATGCCCCAAGTCAATATTGTCAAGAAAACAGAAGTACAGACAGGCGGATTCTCCAAGGAGTCAA<br>TTTTACCAAAAAGAAATTCGGACAAGCTTATTGCTCGTAAAAAAGACTGGGATCCAAAAAAATATGGTGGTTTTGATAGTCCAAC<br>GGTAGCTTATTCAGTCCTAGTGGTTGCTAAGGTGGAAAAAGGGAAATCGAAGAAGTTAAAATCCGTTAAAGAGTTACTAGGGAT<br>CACAATTATGGAAAGAAGTTCCTTTGAAAAAAATCCGATTGACTTTTTAGAAGCTAAAGGATATAAGGAAGTTAAAAAAGACTTA<br>ATCATTAAACTACCTAAATATAGTCTTTTTGAGTTAGAAAACGGTCGTAAACGGATGCTGGCTAGTGCCGGAGAATTACAAAAAG<br>GAAATGAGCTGGCTCTGCCAAGCAAATATGTGAATTTTTTATATTTAGCTAGTCATTATGAAAAGTTGAAGGGTAGTCCAGAAGA<br>TAACGAACAAAAACAATTGTTTGTGGAGCAGCATAAGCATTATTTAGATGAGATTATTGAGCAAATCAGTGAATTTTCTAAGCGT<br>GTTATTTTAGCAGATGCCAATTTAGATAAAGTTCTTAGTGCATATAACAAACATAGAGACAAACCAATACGTGAACAAGCAGAAA<br>ATATTATTCATTTATTTACGTTGACGAATCTTGGAGCTCCCGCTGCTTTTAAATATTTTGATACAACAATTGATCGTAAACGATATA<br>CGTCTACAAAAGAAGTTTTAGATGCCACTCTTATCCATCAATCCATCACTGGTCTTTATGAAACACGCATTGATTTGAGTCAGCTA<br>GGAGGTGACGCTT |
| pMC0_4_56_CDS AcrIIA4 | AATGAACATTAACGACCTCATACGAGAGATTAAGAACAAAGATTACACCGTCAAACTGTCAGGAACTGATAGTAACTCAATCACC<br>CAGCTTATTATCAGGGTAAACAATGATGGGAATGAATATGTGATATCTGAGAGCGAAAACGAGTCTATCGTCGAGAAATTCATTT<br>CCGCTTTTAAGAACGGGTGGAATCAGGAATATGAGGATGAAGAAGAATTTTACAATGACATGCAGACGATCACGTTGAAAAGTG<br>AACTGAACTAAGCTT |
| pMC0_4_60_CDS SpG Cas9 | AATGGATAAGAAATACTCAATAGGCTTAGATATCGGCACAAATAGCGTCGGATGGGCGGTGATCACTGATGAATATAAGGTTCC<br>GTCTAAAAAGTTCAAGGTTCTGGGAAATACAGACCGCCACAGTATCAAAAAAAATCTTATAGGGGCTCTTTTATTTGACAGTGGA<br>GAGACAGCGGAAGCGACTCGTCTGAAACGGACAGCTCGTAGAAGGTATACACGTCGGAAGAATCGTATTTGTTATCTACAGGAG<br>ATTTTTTCAAATGAGATGGCGAAAGTAGATGATAGTTTCTTTCATCGACTTGAAGAGTCTTTTTTGGTGGAAGAAGACAAGAAGC<br>ATGAACGTCATCCTATTTTTGGAAATATAGTAGATGAAGTTGCTTATCATGAGAAATATCCAACTATCTATCATCTGCGAAAAAAA<br>TTGGTAGATTCTACTGATAAAGCGGATTTGCGCTTAATCTATTTGGCCTTAGCGCATATGATTAAGTTTCGTGGTCATTTTTTGATT<br>GAGGGAGATTTAAATCCTGATAATAGTGATGTGGACAAACTATTTATCCAGTTGGTACAAACCTACAATCAATTATTTGAAGAAA<br>ACCCTATTAACGCAAGTGGAGTAGATGCTAAAGCGATTCTTTCTGCACGATTGAGTAAATCAAGACGATTAGAAAATCTCATTGC<br>TCAGCTCCCCGGTGAGAAGAAAAATGGCTTATTTGGGAATCTCATTGCTTTGTCATTGGGTTTGACCCCTAATTTTAAATCAAATT<br>TTGATTTGGCAGAAGATGCTAAATTACAGCTTTCAAAAGATACTTACGATGATGATTTAGATAATTTATTGGCGCAAATTGGAGAT<br>CAATATGCTGATTTGTTTTTGGCAGCTAAGAATTTATCAGATGCTATTTTACTTTCAGATATCCTAAGAGTAAATACTGAAATAACT<br>AAGGCTCCCCTATCAGCTTCAATGATTAAACGCTACGATGAACATCATCAAGACTTGACTCTTTTAAAAGCTTTAGTTCGACAACA<br>ACTTCCAGAAAAGTATAAAGAAATCTTTTTTGATCAATCAAAAAACGGATATGCAGGTTATATTGATGGGGGAGCTAGCCAAGAA<br>GAATTTTATAAATTTATCAAACCAATTTTAGAAAAAATGGATGGTACTGAGGAATTATTGGTGAAACTAAATCGTGAAGATTTGCT<br>GCGCAAGCAACGGACCTTTGACAACGGCTCTATTCCCCATCAAATTCACTTGGGTGAGCTGCATGCTATTTTGAGAAGACAAGAA<br>GACTTTTATCCATTTTTAAAAGACAATCGTGAGAAGATTGAAAAAATCTTGACTTTTCGTATTCCTTATTATGTTGGTCCATTGGCG<br>CGTGGCAATAGTCGTTTTGCATGGATGACTCGGAAGTCTGAAGAAACAATTACCCCATGGAATTTTGAAGAAGTTGTCGATAAAG<br>GTGCTTCAGCTCAATCATTTATTGAACGCATGACAAACTTTGATAAAAATCTTCCAAATGAAAAAGTACTACCAAAACATAGTTTG<br>CTTTATGAGTATTTTACGGTTTATAACGAATTGACAAAGGTCAAATATGTTACTGAAGGAATGCGAAAACCAGCATTTCTTTCAGG<br>TGAACAGAAGAAAGCCATTGTTGATTTACTCTTCAAAACAAATCGAAAAGTAACCGTTAAGCAATTAAAAGAAGATTATTTCAAA<br>AAAATAGAATGTTTTGATAGTGTTGAAATTTCAGGAGTTGAAGATAGATTTAATGCTTCATTAGGTACCTACCATGATTTGCTAAA<br>AATTATTAAAGATAAAGATTTTTTGGATAATGAAGAAAATGAAGATATCTTAGAGGATATTGTTTTAACATTGACCTTATTTGAAG<br>ATAGGGAGATGATTGAGGAAAGACTTAAAACATATGCTCACCTCTTTGATGATAAGGTGATGAAACAGCTTAAACGTCGCCGTTA<br>TACTGGTTGGGGACGTTTGTCTCGAAAATTGATTAATGGTATTAGGGATAAGCAATCTGGCAAAACAATATTAGATTTTTTGAAA<br>TCAGATGGTTTTGCCAATCGCAATTTTATGCAGCTGATCCATGATGATAGTTTGACATTTAAAGAAGACATTCAAAAAGCACAAGT<br>GTCTGGACAAGGCGATAGTTTACATGAACATATTGCAAATTTAGCTGGTAGCCCTGCTATTAAAAAAGGTATTTTACAGACTGTA<br>AAAGTTGTTGATGAATTGGTCAAAGTAATGGGGCGGCATAAGCCAGAAAATATCGTTATTGAAATGGCACGTGAAAATCAGACA<br>ACTCAAAAGGGCCAGAAAAATTCGCGAGAGCGTATGAAACGAATCGAAGAAGGTATCAAAGAATTAGGAAGTCAGATTCTTAA<br>AGAGCATCCTGTTGAAAATACTCAATTGCAAAATGAAAAGCTCTATCTCTATTATCTCCAAAATGGAAGAGACATGTATGTGGAC<br>CAAGAATTAGATATTAATCGTTTAAGTGATTATGATGTCGATCACATTGTTCCACAAAGTTTCCTTAAAGACGATTCAATAGACAA<br>TAAGGTCTTAACGCGTTCTGATAAAAATCGTGGTAAATCGGATAACGTTCCAAGTGAAGAAGTAGTCAAAAAGATGAAAAACTA<br>TTGGAGACAACTTCTAAACGCCAAGTTAATCACTCAACGTAAGTTTGATAATTTAACGAAAGCTGAACGTGGAGGTTTGAGTGAA<br>CTTGATAAAGCTGGTTTTATCAAACGCCAATTGGTTGAAACTCGCCAAATCACTAAGCATGTGGCACAAATTTTGGATAGTCGCAT<br>GAATACTAAATACGATGAAAATGATAAACTTATTCGAGAGGTTAAAGTGATTACCTTAAAATCTAAATTAGTTTCTGACTTCCGAA<br>AAGATTTCCAATTCTATAAAGTACGTGAGATTAACAATTACCATCATGCCCATGATGCGTATCTAAATGCCGTCGTTGGAACTGCT<br>TTGATTAAGAAATATCCAAAACTTGAATCGGAGTTTGTCTATGGTGATTATAAAGTTTATGATGTTCGTAAAATGATTGCTAAGTC<br>TGAGCAAGAAATAGGCAAAGCAACCGCAAAATATTTCTTTTACTCTAATATCATGAACTTCTTCAAAACAGAAATTACACTTGCAA<br>ATGGAGAGATTCGCAAACGCCCTCTAATCGAAACTAATGGGGAAACTGGAGAAATTGTCTGGGATAAAGGGCGAGATTTTGCCA<br>CAGTGCGCAAAGTATTGTCCATGCCCCAAGTCAATATTGTCAAGAAAACAGAAGTACAGACAGGCGGATTCTCCAAGGAGTCAA<br>TTTTACCAAAAAGAAATTCGGACAAGCTTATTGCTCGTAAAAAAGACTGGGATCCAAAAAAATATGGTGGTTTTCTGTGGCCAAC<br>GGTAGCTTATTCAGTCCTAGTGGTTGCTAAGGTGGAAAAAGGGAAATCGAAGAAGTTAAAATCCGTTAAAGAGTTACTAGGGAT<br>CACAATTATGGAAAGAAGTTCCTTTGAAAAAAATCCGATTGACTTTTTAGAAGCTAAAGGATATAAGGAAGTTAAAAAAGACTTA |

| | |
|---|---|
| | ATCATTAAACTACCTAAATATAGTCTTTTTGAGTTAGAAAACGGTCGTAAACGGATGCTGGCTAGTGCCAAGCAATTACAAAAAG<br>GAAATGAGCTGGCTCTGCCAAGCAAATATGTGAATTTTTTATATTTAGCTAGTCATTATGAAAAGTTGAAGGGTAGTCCAGAAGA<br>TAACGAACAAAAACAATTGTTTGTGGAGCAGCATAAGCATTATTTAGATGAGATTATTGAGCAAATCAGTGAATTTTCTAAGCGT<br>GTTATTTTAGCAGATGCCAATTTAGATAAAGTTCTTAGTGCATATAACAAACATAGAGACAAACCAATACGTGAACAAGCAGAAA<br>ATATTATTCATTTATTTACGTTGACGAATCTTGGAGCTCCCGCTGCTTTTAAATATTTTGATACAACAATTGATCGTAAACAATATC<br>GTTCTACAAAAGAAGTTTTAGATGCCACTCTTATCCATCAATCCATCACTGGTCTTTATGAAACACGCATTGATTTGAGTCAGCTAG<br>GAGGTGACGCTT |
| pMC0_TU2-6*_02_Dropout mScarlet-I | TACTAGAGACGGAAAGTGAAACGTGATTTCATGCGTCATTTTGAACATTTTGTAAATCTTATTTAATAATGTGTGCGGCAATTCAC<br>ATTTAATTTATGAATGTTTTCTTAACATCGCGGCAACTCAAGAAACGGCAGGTTCGGATCTTAGCTACTAGAGAAAGAGGAGAAA<br>TACTAGATGGTTTCTAAAGGTGAAGCAGTGATCAAAGAATTTATGCGCTTCAAAGTTCACATGGAAGGTTCTATGAACGGCCACG<br>AATTTGAAATTGAAGGTGAAGGCGAAGGTCGTCCATACGAAGGTACTCAAACCGCAAAACTGAAAGTTACCAAAGGTGGTCCAC<br>TGCCATTTTCTTGGGATATTCTGTCTCCACAATTTATGTACGGTTCTCGTGCATTTATCAAACACCCAGCAGATATTCCAGACTACT<br>ACAAACAATCTTTTCCGGAAGGTTTCAAATGGGAACGTGTTATGAATTTTGAAGATGGTGGTGCAGTTACGGTTACCCAAGATAC<br>CTCTCTGGAAGATGGTACTCTGATCTACAAAGTTAAACTGCGTGGTACTAACTTTCCACCAGATGGTCCAGTTATGCAGAAAAAA<br>ACCATGGGTTGGGAAGCATCTACCGAACGTCTGTACCCAGAAGATGGCGTTCTGAAAGGTGATATCAAAATGGCACTGCGTCTG<br>AAAGATGGCGGTCGTTACCTGGCAGATTTCAAAACCACCTACAAAGCGAAAAAACCAGTTCAAATGCCAGGTGCATACAACGTT<br>GATCGTAAACTGGATATTACCAGCCACAACGAAGATTACACCGTTGTTGAACAATACGAACGTTCTGAAGGCCGTCACTCTACCG<br>GTGGTATGGATGAACTGTACAAATAACCAGGCATCAAATAAAACGAAAGGCTCAGTCGAAAGACTGGGCCTTTCGTTTTATCTGT<br>TGTTTGTCGGTGAACGCTCTCTACTAGAGTCACACTGGCTCACCTTCGGGTGGGCCTTTCTGCGTTTATACGTCTCTAGCT |
| pMC0_TU5-6*_01_Dropout sfGFP | GCTTAGAGACGGAAAGTGAAACGTGATTTCATGCGTCATTTTGAACATTTTGTAAATCTTATTTAATAATGTGTGCGGCAATTCAC<br>ATTTAATTTATGAATGTTTTCTTAACATCGCGGCAACTCAAGAAACGGCAGGTTCGGATCTTAGCTACTAGAGAAAGAGGAGAAA<br>TACTAGATGCGTAAAGGCGAAGAACTTTTCACTGGCGTAGTACCAATTTTAGTGGAGCTGGACGGAGATGTAAATGGTCATAAG<br>TTTTCAGTTCGAGGAGAAGGCGAAGGAGATGCAACTAACGGTAAGCTGACACTAAAGTTTATCTGTACCACGGGTAAACTGCCG<br>GTCCCATGGCCGACACTGGTTACTACACTTACTTATGGTGTACAATGCTTTGCTCGTTATCCTGACCATATGAAGCAGCACGATTT<br>CTTCAAATCGGCCATGCCTGAAGGATACGTGCAAGAACGTACAATTAGCTTTAAAGATGACGGCACCTATAAAACGCGCGCCGA<br>AGTTAAATTCGAGGGTGACACATTGGTTAATCGTATAGAACTTAAGGGCATTGACTTCAAAGAAGATGGCAACATCCTTGGCCAT<br>AAACTGGAATATAATTTTAACTCTCACAATGTCTACATTACGGCGGATAAACAAAAGAATGGCATTAAAGCGAATTTCAAAATCC<br>GCCACAATGTCGAAGACGGCTCGGTTCAACTGGCGGACCATTATCAGCAGAACACCCCAATCGGTGATGGCCCGGTTCTGTTACC<br>TGATAATCATTACCTTTCTACTCAAAGCGTTTTATCTAAAGATCCTAACGAGAAGCGTGATCATATGGTTCTACTGGAATTTGTTAC<br>CGCAGCTGGTATCACGCACGGCATGGATGAGCTGTATAAATGACCAGGCATCAAATAAAACGAAAGGCTCAGTCGAAAGACTG<br>GGCCTTTCGTTTTATCTGTTGTTTGTCGGTGAACGCTCTCTACTAGAGTCACACTGGCTCACCTTCGGGTGGGCCTTTCTGCGTTTA<br>TACGTCTCTAGCT |
| pMC0*_gRNA_Pos4_Ptet | GCTTCAAATCACTGTAATGATCTTAATTCCTAATTTTTGTTGACACTCTATCATTGATAGAGTTATTTTAGTCCAGAGACCGAAAGT<br>GAAACGTGATTTCATGCGTCATTTTGAACATTTTGTAAATCTTATTTAATAATGTGTGCGGCAATTCACATTTAATTTATGAATGTT<br>TTCTTAACATCGCGGCAACTCAAGAAACGGCAGGTTCGGATCTTAGCTACTAGAGAAAGAGGAGAAATACTAGATGCGTAAAGG<br>CGAAGAACTTTTCACTGGCGTAGTACCAATTTTAGTGGAGCTGGACGGAGATGTAAATGGTCATAAGTTTTCAGTTCGAGGAGA<br>AGGCGAAGGAGATGCAACTAACGGTAAGCTGACACTAAAGTTTATCTGTACCACGGGTAAACTGCCGGTCCCATGGCCGACACT<br>GGTTACTACACTTACTTATGGTGTACAATGCTTTGCTCGTTATCCTGACCATATGAAGCAGCACGATTTCTTCAAATCGGCCATGCC<br>TGAAGGATACGTGCAAGAACGTACAATTAGCTTTAAAGATGACGGCACCTATAAAACGCGCGCCGAAGTTAAATTCGAGGGTGA<br>CACATTGGTTAATCGTATAGAACTTAAGGGCATTGACTTCAAAGAAGATGGCAACATCCTTGGCCATAAACTGGAATATAATTTT<br>AACTCTCACAATGTCTACATTACGGCGGATAAACAAAAGAATGGCATTAAAGCGAATTTCAAAATCCGCCACAATGTCGAAGACG<br>GCTCGGTTCAACTGGCGGACCATTATCAGCAGAACACCCCAATCGGTGATGGCCCGGTTCTGTTACCTGATAATCATTACCTTTCT<br>ACTCAAAGCGTTTTATCTAAAGATCCTAACGAGAAGCGTGATCATATGGTTCTACTGGAATTTGTTACCGCAGCTGGTATCACGCA<br>CGGCATGGATGAGCTGTATAAATAACCAGGCATCAAATAAAACGAAAGGCTCAGTCGAAAGACTGGGCCTTTCGTTTTATCTGTT<br>GTTTGTCGGTGAACGCTCTCTACTAGAGTCACACTGGCTCACCTTCGGGTGGGCCTTTCTGCGTTTATAGGTCTCAGTTTCAGAGC<br>TATGCTGGAAACAGCATAGCAAGTTGAAATAAGGCTAGTCCGTTATCAACTTGAAAAAGTGGCACCGAGTCGGTGCTTTTTTCTC<br>GGTACCAAATTCCAGAAAAGAGGCCTCCCGAAAGGGGGGCCTTTTTTCGTTTTGGTCCACCCTGCACGATAACACTGAGGTA |
| pMC0*_gRNA_Pos5a_Ptet | GGTACAAATCACTGTAATGATCTTAATTCCTAATTTTTGTTGACACTCTATCATTGATAGAGTTATTTTAGTCCAGAGACCGAAAGT<br>GAAACGTGATTTCATGCGTCATTTTGAACATTTTGTAAATCTTATTTAATAATGTGTGCGGCAATTCACATTTAATTTATGAATGTT<br>TTCTTAACATCGCGGCAACTCAAGAAACGGCAGGTTCGGATCTTAGCTACTAGAGAAAGAGGAGAAATACTAGATGCGTAAAGG<br>CGAAGAACTTTTCACTGGCGTAGTACCAATTTTAGTGGAGCTGGACGGAGATGTAAATGGTCATAAGTTTTCAGTTCGAGGAGA<br>AGGCGAAGGAGATGCAACTAACGGTAAGCTGACACTAAAGTTTATCTGTACCACGGGTAAACTGCCGGTCCCATGGCCGACACT<br>GGTTACTACACTTACTTATGGTGTACAATGCTTTGCTCGTTATCCTGACCATATGAAGCAGCACGATTTCTTCAAATCGGCCATGCC<br>TGAAGGATACGTGCAAGAACGTACAATTAGCTTTAAAGATGACGGCACCTATAAAACGCGCGCCGAAGTTAAATTCGAGGGTGA<br>CACATTGGTTAATCGTATAGAACTTAAGGGCATTGACTTCAAAGAAGATGGCAACATCCTTGGCCATAAACTGGAATATAATTTT<br>AACTCTCACAATGTCTACATTACGGCGGATAAACAAAAGAATGGCATTAAAGCGAATTTCAAAATCCGCCACAATGTCGAAGACG<br>GCTCGGTTCAACTGGCGGACCATTATCAGCAGAACACCCCAATCGGTGATGGCCCGGTTCTGTTACCTGATAATCATTACCTTTCT<br>ACTCAAAGCGTTTTATCTAAAGATCCTAACGAGAAGCGTGATCATATGGTTCTACTGGAATTTGTTACCGCAGCTGGTATCACGCA<br>CGGCATGGATGAGCTGTATAAATAACCAGGCATCAAATAAAACGAAAGGCTCAGTCGAAAGACTGGGCCTTTCGTTTTATCTGTT<br>GTTTGTCGGTGAACGCTCTCTACTAGAGTCACACTGGCTCACCTTCGGGTGGGCCTTTCTGCGTTTATAGGTCTCAGTTTCAGAGC<br>TATGCTGGAAACAGCATAGCAAGTTGAAATAAGGCTAGTCCGTTATCAACTTGAAAAAGTGGCACCGAGTCGGTGCTTTTTTCTC<br>GGTACCAAATTCCAGAAAAGAGGCCTCCCGAAAGGGGGGCCTTTTTTCGTTTTGGTCCACCCTGCACGATAACACTGAGTAA |
| pMC0*_gRNA_Pos5b_Ptet | GTAACAAATCACTGTAATGATCTTAATTCCTAATTTTTGTTGACACTCTATCATTGATAGAGTTATTTTAGTCCAGAGACCGAAAGT<br>GAAACGTGATTTCATGCGTCATTTTGAACATTTTGTAAATCTTATTTAATAATGTGTGCGGCAATTCACATTTAATTTATGAATGTT<br>TTCTTAACATCGCGGCAACTCAAGAAACGGCAGGTTCGGATCTTAGCTACTAGAGAAAGAGGAGAAATACTAGATGCGTAAAGG<br>CGAAGAACTTTTCACTGGCGTAGTACCAATTTTAGTGGAGCTGGACGGAGATGTAAATGGTCATAAGTTTTCAGTTCGAGGAGA<br>AGGCGAAGGAGATGCAACTAACGGTAAGCTGACACTAAAGTTTATCTGTACCACGGGTAAACTGCCGGTCCCATGGCCGACACT |

| | |
|---|---|
| | GGTTACTACACTTACTTATGGTGTACAATGCTTTGCTCGTTATCCTGACCATATGAAGCAGCACGATTTCTTCAAATCGGCCATGCC<br>TGAAGGATACGTGCAAGAACGTACAATTAGCTTTAAAGATGACGGCACCTATAAAACGCGCGCCGAAGTTAAATTCGAGGGTGA<br>CACATTGGTTAATCGTATAGAACTTAAGGGCATTGACTTCAAAGAAGATGGCAACATCCTTGGCCATAAACTGGAATATAATTTT<br>AACTCTCACAATGTCTACATTACGGCGGATAAACAAAAGAATGGCATTAAAGCGAATTTCAAAATCCGCCACAATGTCGAAGACG<br>GCTCGGTTCAACTGGCGGACCATTATCAGCAGAACACCCCAATCGGTGATGGCCCGGTTCTGTTACCTGATAATCATTACCTTTCT<br>ACTCAAAGCGTTTTATCTAAAGATCCTAACGAGAAGCGTGATCATATGGTTCTACTGGAATTTGTTACCGCAGCTGGTATCACGCA<br>CGGCATGGATGAGCTGTATAAATAACCAGGCATCAAATAAAACGAAAGGCTCAGTCGAAAGACTGGGCCTTTCGTTTTATCTGTT<br>GTTTGTCGGTGAACGCTCTCTACTAGAGTCACACTGGCTCACCTTCGGGTGGGCCTTTCTGCGTTTATAGGTCTCAGTTTCAGAGC<br>TATGCTGGAAACAGCATAGCAAGTTGAAATAAGGCTAGTCCGTTATCAACTTGAAAAAGTGGCACCGAGTCGGTGCTTTTTTCTC<br>GGTACCAAATTCCAGAAAAGAGGCCTCCCGAAAGGGGGGCCTTTTTTCGTTTTGGTCCACCCTGCACGATAACACTGAAGCT |

*Supplementary Table S2: Assembly of plasmids used in this study. Parts were assembled in the framework of the Marburg Collection* (Stukenberg *et al.*, 2021). *Short nomenclature is provided in the table, e.g. 2_43 = pMC0_**2_43**_Ptac. Parts and plasmids written in bold letters were assembled in this project and all remaining parts are available as genetic parts in the Marburg Collection. Part sequences for new parts are provided in Supplementary Table S1.*

| Plasmid | Parts and plasmid used for assembly | Description |
|---|---|---|
| pST_025 | 1*-6* Linker, 8*_06, 7*_01 | Control plasmid |
| pST_032 | pDS_120, **2_43**, 3_07, **4_45**, 5_08 | Level 1, $P_{tac}$ Tfox |
| pST_033 | pDS_191, **2_42**, 3_04, **4_25**, 5a_07, 5b_02 | Level 1, $P_{tet}$ Cas9, RBS = B0031 |
| pST_034 | pDS_191, **2_42**, 3_02, **4_25**, 5a_07, 5b_02 | Level 1, $P_{tet}$ Cas9, RBS = B0029 |
| pST_035 | pDS_191, **2_42**, 3_06, **4_25**, 5a_07, 5b_02 | Level 1, $P_{tet}$ Cas9, RBS = B0033 |
| pST_040 | **pST_032**, 8*_06, 7*_01, **1*-6*_03 Dropout** | Level 2 Dropout plasmid, contains $P_{tac}$ Tfox and Dropout for remaining positions |
| pST_084 | pDS_187, 2_08, 3_04, **4_56**, 5_05 | Level 1, J23106 AcrIIA4, RBS = B0031 |
| pST_085 | pDS_187, 2_08, 3_05, **4_56**, 5_05 | Level 1, J23106 AcrIIA4, RBS = B0032 |
| pST_086 | pDS_187, 2_08, 3_07, **4_56**, 5_05 | Level 1, J23106 AcrIIA4, RBS = B0034 |
| pST_087 | pDS_187, 2_08, 3_03, **4_56**, 5_05 | Level 1, J23106 AcrIIA4, RBS = B0030 |
| pST_107 | **pST_040**, **gRNA Pos 4 ($P_{tet}$)**, TU5-6*_EL_04, **pST_033**, **pST_084** | Level 2, NT-CRISPR, Cas9 RBS = B0031, AcrIIA4 RBS = B0031 |
| pST_108 | **pST_040**, **gRNA Pos 4 ($P_{tet}$)**, TU5-6*_EL_04, **pST_033**, **pST_085** | Level 2, NT-CRISPR, Cas9 RBS = B0031, AcrIIA4 RBS = B0032 |
| pST_109 | **pST_040**, **gRNA Pos 4 ($P_{tet}$)**, TU5-6*_EL_04, **pST_033**, **pST_086** | Level 2, NT-CRISPR, Cas9 RBS = B0031, AcrIIA4 RBS = B0034 |
| pST_110 | **pST_040**, **gRNA Pos 4 ($P_{tet}$)**, TU5-6*_EL_04, **pST_033**, **pST_087** | Level 2, NT-CRISPR, Cas9 RBS = B0031, AcrIIA4 RBS = B0030 |
| pST_111 | **pST_040**, **gRNA Pos 4 ($P_{tet}$)**, TU5-6*_EL_04, **pST_034**, **pST_084** | Level 2, NT-CRISPR, Cas9 RBS = B0029, AcrIIA4 RBS = B0031 |
| pST_112 | **pST_040**, **gRNA Pos 4 ($P_{tet}$)**, TU5-6*_EL_04, **pST_034**, **pST_085** | Level 2, NT-CRISPR, Cas9 RBS = B0029, AcrIIA4 RBS = B0032 |
| pST_113 | **pST_040**, **gRNA Pos 4 ($P_{tet}$)**, TU5-6*_EL_04, **pST_034**, **pST_086** | Level 2, NT-CRISPR, Cas9 RBS = B0029, AcrIIA4 RBS = B0034 |
| pST_114 | **pST_040**, **gRNA Pos 4 ($P_{tet}$)**, TU5-6*_EL_04, **pST_034**, **pST_087** | Level 2, NT-CRISPR, Cas9 RBS = B0029, AcrIIA4 RBS = B0030 |
| pST_115 | **pST_040**, **gRNA Pos 4 ($P_{tet}$)**, TU5-6*_EL_04, **pST_035**, **pST_084** | Level 2, NT-CRISPR, Cas9 RBS = B0033, AcrIIA4 RBS = B0031 |

| | | |
|---|---|---|
| pST_116 | **pST_040**, **gRNA Pos 4 ($P_{tet}$)**, TU5-6*_EL_04, **pST_035**, **pST_085** | Level 2, NT-CRISPR, Cas9 RBS = B0033, AcrIIA4 RBS = B0032 |
| pST_117 | **pST_040**, **gRNA Pos 4 ($P_{tet}$)**, TU5-6*_EL_04, **pST_035**, **pST_086** | Level 2, NT-CRISPR, Cas9 RBS = B0033, AcrIIA4 RBS = B0034 |
| pST_118 | **pST_040**, **gRNA Pos 4 ($P_{tet}$)**, TU5-6*_EL_04, **pST_035**, **pST_087** | Level 2, NT-CRISPR, Cas9 RBS = B0033, AcrIIA4 RBS = B0030 |
| pST_119 | **pST_040**, **pST_035**, **pST_085**, **1*-6*_09 Dropout** | NT-CRISPR plasmid for multiple gRNAs, Cas9 RBS = B0033, AcrIIA4 RBS = B0032, Dropout for positions 4 and 5 |
| pST_133 | pDS_120, 2_13, 3_03, 4_12, 5_03 | Level 1, strong constitutive mScarlet-I, used as template to construct tDNA for integration |
| pST_136 | pDS_191, 2_42, 3_06, 4_60, 5a_07, 5b_02 | Level 1, $P_{tet}$ SpG Cas9, RBS = B0033 |
| pST_137 | **pST_119**, **gRNA Pos4 ($P_{tet}$, flrB)**, **gRNA Pos5a ($P_{tet}$, vnp1)**, **gRNA Pos5b ($P_{tet}$, vnp2)** | Level 2, NT-CRISPR plasmid with 3 gRNAs (flrB, vnp1, vnp2) |
| pST_138 | **gRNA Pos4 ($P_{tet}$, xds)**, **gRNA Pos5a ($P_{tet}$, pilA)**, **gRNA Pos5b ($P_{tet}$, galE)** | Level 2, NT-CRISPR plasmid with 3 gRNAs (flrB, vnp1, vnp2) |
| pST_140 | **pST_040**, **gRNA Pos 4 ($P_{tet}$)**, TU5-6*_EL_04, **pST_136**, **pST_085** | Level 2, NT-CRISPR with SpG Cas9 |

*Supplementary Table S3: Oligonucleotides used to assemble gRNA sequences. All sequences written as 5' -> 3'.*

| Target | gRNA sequence | forward oligonucleotide | | reverse oligonucleotide | |
|---|---|---|---|---|---|
| wbfF | TTAGCCAAGATCAGTCACGT | oDS_200_A_gRNA_wbf 1_fwd | GTCCTTAGCCAAGATCAGTCACGT | oDS_201_A_gRNA_wbf 1_rev | AAACACGTGACTGATCTTGGCTAA |
| xds | ATATTCCGTAATCAAAACTT | oDS_295_A_gRNA_xds_fwd | GTCCATATTCCGTAATCAAAACTT | oDS_296_A_gRNA_xds_rev | AAACAAGTTTTGATTACGGAATAT |
| flrA (flagella) | AAGGATTAAAACGTTAGCTT | oDS_299_A_gRNA_flrA_fwd | GTCCAAGGATTAAAACGTTAGCTT | oDS_300_A_gRNA_flrA_rev | AAACAAGCTAACGTTTTAATCCTT |
| pilA (pili) | CTGATTTCTTTACGTAGTTT | oDS_309_A_gRNA_pilA_fwd | GTCCCTGATTTCTTTACGTAGTTT | oDS_310_A_gRNA_pilA_rev | AAACAAACTACGTAAAGAAATCAG |
| galE | ATGTGTTTACTACTACCGTA | oDS_517_A_gRNA_galE_fwd | GTCCATGTGTTTACTACTACCGTA | oDS_518_A_gRNA_galE_rev | AAACTACGGTAGTAGTAAACACAT |
| vnp1 | ATGGAAACAGACCTACCCAG | oDS_551_A_gRNA_vnp1_fwd | GTCCATGGAAACAGACCTACCCAG | oDS_552_A_gRNA_vnp1_rev | AAACCTGGGTAGGTCTGTTTCCAT |
| vnp2 | TACTCACTGGAGCAGCAGCA | oDS_563_A_gRNA_vnp2_fwd | GTCCTACTCACTGGAGCAGCAGCA | oDS_564_A_gRNA_vnp2_rev | AAACTGCTGCTGCTCCAGTGAGTA |
| rpoS (indirect) | AGTTCTGTCGACACGCCAAT | oDS_380_gRNA_rpoS_fwd | GTCCAGTTCTGTCGACACGCCAAT | oDS_381_gRNA_rpoS_rev | AAACATTGGCGTGTCGACAGAACT |
| hisG | TTGAAAAAAATGATGGAGTAA | oDS_435_A_gRNA_hisG_fwd | GTCCTTGAAAAAAATGATGGAGTAA | oDS_436_A_gRNA_hisG_rev | AAACTTACTCCATCATTTTTTCAA |
| rplS (indirect) | CACCTGGTGCAAATTTAGGT | oDS_448_A_gRNA_rplS_ind_fwd | GTCCCACCTGGTGCAAATTTAGGT | oDS_449_A_gRNA_rplS_ind_rev | AAACACCTAAATTTGCACCAGGTG |
| rpsB | TCGTAGAAGCTGAATAATAG | oDS_459_A_gRNA_rpsB_fwd | GTCCTCGTAGAAGCTGAATAATAG | oDS_460_A_gRNA_rpsB_rev | AAACCTATTATTCAGCTTCTACGA |
| intergenic integration | AGTCGCTTATGGCGTGAAAG | oDS_473_A_gRNA_int9_fwd | GTCCAGTCGCTTATGGCGTGAAAG | oDS_474_A_gRNA_int9_rev | AAACCTTTCACGCCATAAGCGACT |
| malQ | TGGCGTTTAGAAACAGAGCA | oDS_399_A_gRNA_malQ_fwd | GTCCTGGCGTTTAGAAACAGAGCA | oDS_400_A_gRNA_malQ_rev | AAACTGCTCTGTTTCTAAACGCCA |
| glpK | ACTCAGATTTATCCGCAAGC | oDS_596_A_gRNA_glpK_fwd | GTCCACTCAGATTTATCCGCAAGC | oDS_597_A_gRNA_glpK_rev | AAACGCTTGCGGATAAATCTGAGT |

| | | | | | |
|---|---|---|---|---|---|
| wbfF (PAM = NGA) | GTAAGAAACCCATTCCGCTT | oDS_617_A_gRNA_wbf NGA_fwd | GTCCGTAAGAAACCCATTCCGCTT | oDS_618_A_gRNA_wbf NGA_rev | AAACAAGCGGAATGGGTTTCTTAC |
| wbfF (PAM = NGC) | GATTGGTAATCGTGACAGCG | oDS_619_A_gRNA_wbf NGC_fwd | GTCCGATTGGTAATCGTGACAGCG | oDS_620_A_gRNA_wbf NGC_rev | AAACCGCTGTCACGATTACCAATC |
| wbfF (PAM = NGT) | TGGATATCCAAGTATCAGGG | oDS_621_A_gRNA_wbf NGT_fwd | GTCCTGGATATCCAAGTATCAGGG | oDS_622_A_gRNA_wbf NGT_rev | AAACCCCTGATACTTGGATATCCA |
| wbfF (PAM = NGC) (2) | ACGCGGGAATGGCTTCAGAT | oDS_646_A_gRNA_wbf_NGC_2_fwd | GTCCACGCGGGAATGGCTTCAGAT | oDS_647_A_gRNA_wbf_NGC_2_rev | AAACATCTGAAGCCATTCCCGCGT |
| wbfF (PAM = NGA) (2) | ATACGTAAACTTACTGACCG | oDS_648_A_gRNA_wbf_NGA_2_fwd | GTCCATACGTAAACTTACTGACCG | oDS_649_A_gRNA_wbf_NGA_2_rev | AAACCGGTCAGTAAGTTTACGTAT |
| rpoS (PAM = NGT) | TGTCGATTAGTCATCTTCGA | oDS_623_A_gRNA_rpoS NGT_fwd | GTCCTGTCGATTAGTCATCTTCGA | oDS_624_A_gRNA_rpoS NGT_rev | AAACTCGAAGATGACTAATCGACA |

*Supplementary Table S4: Oligonucleotides used for the construction of tDNA template plasmids. All sequences written as 5' -> 3'.*

| **Plasmid** | **forward oligonucleotide** | | **reverse oligonucleotide** | | **PCR template** |
|---|---|---|---|---|---|
| wbfF tDNA template | oDS_193_GA_dwbf_U_fwd | TAGCAACTGTTTTAGCGCTGAGC | oDS_194_GA_dwbf_U_rev | TATACATCAATTGCTTTTATCATCATACTCATTCATTAAAG | V. natriegens DNA |
| | oDS_195_GA_dwbf_D_fwd | TGATGATAAAAGCAATTGATGTATAAGCGTCATTTATTCG | oDS_196_GA_dwbf_D_rev | GTGTTCCTGTCGATAAGTATTGATC | V. natriegens DNA |
| | oDS_197_GA_dwbf_V_rev | ACGCTCAGCGCTAAAACAGTTGCTAGGCCGCGAATTCCAGAAATC | oDS_198_GA_dwbf_V_fwd | GATCAATACTTATCGACAGGAACACTAGTAGCGGCCGCTGCAGTC | pMC_V*_03 |
| xds tDNA template | oDS_531_GA_xds_U_fwd | TCGGTTACGTTTTTCAGGAGC | oDS_532_GA_xds_U_rev | AAACACTTATGTCCCATGTTTTATAGATTGAATGTTTAATATCG | V. natriegens DNA |
| | oDS_533_GA_xds_D_fwd | TATAAAACATGGGACATAAGTGTTTATTGAAGTGAAGG | oDS_534_GA_xds_D_rev | TGCCGATTTTCGTCCCGATC | V. natriegens DNA |
| | oDS_536_GA_xds_V_fwd | GTGATGATCGGGACGAAAATCGGCATAGTAGCGGCCGCTGCAGTC | oDS_535_GA_xds_V_rev | CTTGGCTCCTGAAAAACGTAACCGAGGCCGCGAATTCCAGAAATC | pMC_V_01 |
| flagella tDNA Template | oDS_355_GA_flagella_U_fwd | ACTTACAGAATGAGCGTAAATGG | oDS_356_GA_flagella_U_rev | AGTCAGTGTTTTAAATATGTGCTCCAATAAGATGAAGG | V. natriegens DNA |
| | oDS_357_GA_flagella_D_fwd | GGAGCACATATTTAAAACACTGACTTTATGATTCAGTGG | oDS_358_GA_flagella_D_rev | TGTCAACGGCACTTTGATGG | V. natriegens DNA |
| | oDS_360_GA_flagella_V_fwd | GATTACCATCAAAGTGCCGTTGACATAGTAGCGGCCGCTGCAGTC | oDS_359_GA_flagella_V_rev | TCCCATTTACGCTCATTCTGTAAGTGGCCGCGAATTCCAGAAATC | pMC_V_01 |
| pili tDNA template | oDS_497_GA_pili_U_fwd | CAGTTGCTCGAAAATCTCAACC | oDS_498_GA_pili_U_rev | GAAGGAAGAGAGATTTCACTTTGAACTAGCAGCCATTTAG | V. natriegens DNA |
| | oDS_499_GA_pili_D_fwd | TTCAAAGTGAAATCTCTCTTCCTTCTCATGAGATAG | oDS_500_GA_pili_D_rev | TGGTTGGATCTATAAAACAAGCAAATC | V. natriegens DNA |
| | oDS_502_GA_pili_V_fwd | TTTGCTTGTTTTATAGATCCAACCATAGTAGCGGCCGCTGCAGTC | oDS_501_GA_pili_V_rev | TATGGTTGAGATTTTCGAGCAACTGGGCCGCGAATTCCAGAAATC | pMC_V_01 |
| galE tDNA template | oDS_507_GA_galE_U_fwd | GCAAGGCGCACGATTAAACC | oDS_508_GA_galE_U_rev | CGGAGAAGTAAAATTTTCATCCGGTGCAGTGCG | V. natriegens DNA |
| | oDS_509_GA_galE_D_fwd | ACCGGATGAAAATTTTACTTCTCCGCTGTTGCTG | oDS_510_GA_galE_D_rev | ATTTTTTATTGGAACTTGACTGGTTAG | V. natriegens DNA |
| | oDS_512_GA_galE_V_fwd | AACCAGTCAAGTTCCAATAAAAATTAGTAGCGGCCGCTGCAGTC | oDS_511_GA_galE_V_rev | GATTTGGTTTAATCGTGCGCCTTGCGGCCGCGAATTCCAGAAATC | pMC_V_01 |
| | oDS_541_GA_vnp1_U_fwd | TCGGCACACAAATTGGTACG | oDS_542_GA_vnp1_U_rev | AAAAACCACACAAACAAATCTACACTCACGTTATCG | V. natriegens DNA |

| | | | | | |
|---|---|---|---|---|---|
| vnp1 tDNA template | oDS_543_GA_vnp1_D_fwd | GTGTAGATTTGTTTGTGTGGTTTTTACATCTGAGTG | oDS_544_GA_vnp1_D_rev | ACCACAACACTCAATTTGGACG | V. natriegens DNA |
| | oDS_546_GA_vnp1_V_fwd | CGGCGTCCAAATTGAGTGTTGTGGTTAGTAGCGGCCGCTGCAGTC | oDS_545_GA_vnp1_V_rev | AATAGCGTACCAATTTGTGTGCCGAGGCCGCGAATTCCAGAAATC | pMC_V_01 |
| vnp2 tDNA template | oDS_553_GA_vnp2_U_fwd | CAAATCCTTCGGACAGCAGAG | oDS_554_GA_vnp2_U_rev | TTCGTAGTCAAAGGAAGAAAAAAGTGGGCTGATTATC | V. natriegens DNA |
| | oDS_555_GA_vnp2_D_fwd | ACTTTTTTCTTCCTTTGACTACGAAACTTGGGGAAC | oDS_556_GA_vnp2_D_rev | GCAGAGCAACACGCAGCTC | V. natriegens DNA |
| | oDS_558_GA_vnp2_V_fwd | GACAAAGAGCTGCGTGTTGCTCTGCTAGTAGCGGCCGCTGCAGTC | oDS_557_GA_vnp2_V_rev | TCTACTCTGCTGTCCGAAGGATTTGGGCCGCGAATTCCAGAAATC | pMC_V_01 |
| rpoS mScarlet-I tDNA template (indirect) | oDS_366_GA_rpoS_U_fwd | AAGCGCCATTTTGAGTCTGC | oDS_367_GA_rpoS_U_Mut_rev | TCACCACCAATAGGCGTGTCGACAG | V. natriegens DNA |
| | oDS_368_GA_rpoS_U_Mut_fwd | CTGTCGACACGCCTATTGGTGGTG | oDS_369_GA_rpoS_U_rev | TGATCACTGCTTCACCTTTAGAAACGTCATCTTCGACGTTAAACAAG | V. natriegens DNA |
| | oDS_370_GA_rpoS_D_fwd | CGGTGGTATGGATGAACTGTACAAATAATCGACATACATAAAGAGAAAAGGC | oDS_371_GA_rpoS_D_rev | ACTGATCGAGACGCCTATCG | V. natriegens DNA |
| | oDS_373_GA_rpoS_V_fwd | TGCAACGATAGGCGTCTCGATCAGTTAGTAGCGGCCGCTGCAGTC | oDS_372_GA_rpoS_V_rev | TTGCCGCAGACTCAAAATGGCGCTTGGCCGCGAATTCCAGAAATC | pMC_V_01 |
| | oDS_374_GA_mScarlet-I_fwd | GTTTCTAAAGGTGAAGCAGTGATC | oDS_375_GA_mScarlet-I_rev | TTTGTACAGTTCATCCATACCAC | pMC0_4_12_CDSmScarlet-I-I (Vn) |
| rpoS mScarlet-I tDNA template (direct) | oDS_366_GA_rpoS_U_fwd | AAGCGCCATTTTGAGTCTGC | oDS_369_GA_rpoS_U_rev | TGATCACTGCTTCACCTTTAGAAACGTCATCTTCGACGTTAAACAAG | V. natriegens DNA |
| | oDS_370_GA_rpoS_D_fwd | CGGTGGTATGGATGAACTGTACAAATAATCGACATACATAAAGAGAAAAGGC | oDS_371_GA_rpoS_D_rev | ACTGATCGAGACGCCTATCG | V. natriegens DNA |
| | oDS_373_GA_rpoS_V_fwd | TGCAACGATAGGCGTCTCGATCAGTTAGTAGCGGCCGCTGCAGTC | oDS_372_GA_rpoS_V_rev | TTGCCGCAGACTCAAAATGGCGCTTGGCCGCGAATTCCAGAAATC | pMC_V_01 |
| | oDS_374_GA_mScarlet-I_fwd | GTTTCTAAAGGTGAAGCAGTGATC | oDS_375_GA_mScarlet-I_rev | TTTGTACAGTTCATCCATACCAC | pMC0_4_12_CDSmScarlet-I-I (Vn) |
| tDNA template hisG-mScarlet-I | oDS_426_GA_hisG_U_fwd | TCACGCCATCTTCACCATCAAAG | oDS_427_GA_hisG_U_rev | TGATCACTGCTTCACCTTTAGAAACCTCCATCATTTTTTCAATTGGTAGC | V. natriegens DNA |
| | oDS_428_GA_hisG_D_fwd | CGGTGGTATGGATGAACTGTACAAATAAGGGTCATGAGAACAGTTGTTTG | oDS_429_GA_hisG_D_rev | CCTTGCTCATCAAGATTCACG | V. natriegens DNA |
| | oDS_431_GA_hisG_V_fwd | TTGCCGTGAATCTTGATGAGCAAGGTAGTAGCGGCCGCTGCAGTC | oDS_430_GA_hisG_V_rev | TACTTTGATGGTGAAGATGGCGTGAGGCCGCGAATTCCAGAAATC | pMC_V_01 |
| | oDS_374_GA_mScarlet-I_fwd | GTTTCTAAAGGTGAAGCAGTGATC | oDS_375_GA_mScarlet-I_rev | TTTGTACAGTTCATCCATACCAC | pMC0_4_12_CDSmScarlet-I-I (Vn) |
| tDNA template rplS-mScarlet-I (direct) | oDS_437_GA_rplS_U_fwd | GATGATGGGTGAGATTAAAGATC | oDS_440_GA_rplS_U_rev | TGATCACTGCTTCACCTTTAGAAACCTTCTTAGCAAGTTTCTCTTTGATAC | V. natriegens DNA |
| | oDS_441_GA_rplS_D_fwd | CGGTGGTATGGATGAACTGTACAAATAATGCTATAACTAGCGTTCTCATTAAAC | oDS_442_GA_rplS_D_rev | TTATTGTTTTTCTTGGTGTGGTCG | V. natriegens DNA |
| | oDS_444_GA_rplS_V_fwd | ACGACCACACCAAGAAAAACAATAATAGTAGCGGCCGCTGCAGTC | oDS_443_GA_rplS_V_rev | AAGATCTTTAATCTCACCCATCATCGGCCGCGAATTCCAGAAATC | pMC_V_11 |
| | oDS_374_GA_mScarlet-I_fwd | GTTTCTAAAGGTGAAGCAGTGATC | oDS_375_GA_mScarlet-I_rev | TTTGTACAGTTCATCCATACCAC | pMC0_4_12_CDSmScarlet-I-I (Vn) |

| | | | | | |
|---|---|---|---|---|---|
| tDNA template rplS-mScarlet-I (indirect) | oDS_439_GA_rplS_U_Mut_fwd | ATCAGACCTACCAAAATTTGCACCAGGTGAC | oDS_442_GA_rplS_D_rev | TTATTGTTTTTCTTGGTGTGGTCG | tDNA template rplS-mScarlet-I (direct) |
| | oDS_444_GA_rplS_V_fwd | ACGACCACACCAAGAAAAACAATAATAGTAGCGGCCGCTGCAGTC | oDS_438_GA_rplS_U_Mut_rev | TGGTGCAAATTTTGGTAGGTCTGATTTCATTTG | tDNA template rplS-mScarlet-I (direct) |
| tDNA template rpsB-mScarlet-I | oDS_450_GA_rpsB_U_fwd | GATAAAACTCCTTCATCATCATTTCG | oDS_451_GA_rpsB_U_rev | TGATCACTGCTTCACCTTTAGAAACTTCAGCTTCTACGAAACCGTC | V. natriegens DNA |
| | oDS_452_GA_rpsB_D_fwd | CGGTGGTATGGATGAACTGTACAAATAATAGCGGCTCTAGTCACAC | oDS_453_GA_rpsB_D_rev | ATACCAGTGTTTTCTGCCGTTAG | V. natriegens DNA |
| | oDS_455_GA_rpsB_V_fwd | AGCTAACGGCAGAAAACACTGGTATTAGTAGCGGCCGCTGCAGTC | oDS_454_GA_rpsB_V_rev | GAAATGATGATGAAGGAGTTTTATCGGCCGCGAATTCCAGAAATC | pMC_V_11 |
| | oDS_374_GA_mScarlet-I_fwd | GTTTCTAAAGGTGAAGCAGTGATC | oDS_375_GA_mScarlet-I_rev | TTTGTACAGTTCATCCATACCAC | pMC0_4_12_CDSmScarlet-I-I (Vn) |
| int9 mScarlet-I integration tDNA template | oDS_461_GA_int9_U_fwd | TTTGTTTCGCCAGAGTATGGAG | oDS_462_GA_int9_U_rev | CGTCTTTTTTCGTTTTGGTCCTGTTAAAGGGGCACGTTTTTCTATTCTTTAAC | V. natriegens DNA |
| | oDS_463_GA_int9_D_fwd | GCCTTTCTGCGTTTATACGCTTACTCACGCCATAAGCGACTAAGAAAC | oDS_464_GA_int9_D_rev | CTCCATGTTTATGTCTAACACGG | V. natriegens DNA |
| | oDS_466_GA_int9_V_fwd | TGCCGTGTTAGACATAAACATGGAGTAGTAGCGGCCGCTGCAGTC | oDS_465_GA_int9_V_rev | ACACTCCATACTCTGGCGAAACAAAGGCCGCGAATTCCAGAAATC | pMC_V_11 |
| | oDS_467_GA_integration_insert_fwd | AACAGGACCAAAACGAAAAAAGACG | oDS_468_GA_integration_insert_rev | AGTAAGCGTATAAACGCAGAAAGG | pST_133 |
| malQ 298C>T tDNA Template | oDS_391_GA_malQ_U_fwd | CAAATATGGCACTAAGTGTTCAGC | oDS_392_GA_malQ_U_rev | TTAGAAACAGAGTAAGGAGAGGTACTTGAAGGC | V. natriegens DNA |
| | oDS_393_GA_malQ_D_fwd | GTACCTCTCCTTACTCTGTTTCTAAACGCCAGC | oDS_394_GA_malQ_D_rev | AAAGCTTTGTTGCGATTCAGATC | V. natriegens DNA |
| | oDS_396_GA_malQ_V_fwd | GTGATCTGAATCGCAACAAAGCTTTTAGTAGCGGCCGCTGCAGTC | oDS_395_malQ_V_rev | GGCTGAACACTTAGTGCCATATTTGGGCCGCGAATTCCAGAAATC | pMC_V_01 |
| glpK 76C>T tDNA template | oDS_587_GA_glpK_U_fwd | CCAAGAATGGCGTTTGCTGC | oDS_588_GA_glpK_U_rev | CCCAACCTGCTTACGGATAAATCTGAGTAAATTC | V. natriegens DNA |
| | oDS_589_GA_glpK_D_fwd | CAGATTTATCCGTAAGCAGGTTGGGTTGAG | oDS_590_GA_glpK_D_rev | ATTCAAAACCGGTTTTTATCTCCG | V. natriegens DNA |
| | oDS_592_GA_glpK_V_fwd | CCGGAGATAAAAACCGGTTTTGAATTAGTAGCGGCCGCTGCAGTC | oDS_591_GA_glpK_V_rev | CCGTGGCAGCAAACGCCATTCTTGGGGCCGCGAATTCCAGAAATC | pMC_V_01 |
| glpK 76C>T, G81>C tDNA template | oDS_587_GA_glpK_U_fwd | CCAAGAATGGCGTTTGCTGC | oDS_644_GA_glpK C:C_U_rev | CCCAACCTGCTTAGGGATAAATCTGAGTAAATTC | V. natriegens DNA |
| | oDS_645_GA_glpK C:C_U_fwd | CAGATTTATCCCTAAGCAGGTTGGGTTGAG | oDS_590_GA_glpK_D_rev | ATTCAAAACCGGTTTTTATCTCCG | V. natriegens DNA |
| | oDS_592_GA_glpK_V_fwd | CCGGAGATAAAAACCGGTTTTGAATTAGTAGCGGCCGCTGCAGTC | oDS_591_GA_glpK_V_rev | CCGTGGCAGCAAACGCCATTCTTGGGGCCGCGAATTCCAGAAATC | pMC_V_01 |
| araA C123>T tDNA template | oDS_565_GA_araA_U_fwd | ATAGTTACGCCAGATAGGGCAG | oDS_566_GA_araA_U_rev | TCTTTGCCACTTACTCCAATACTTTAGGACC | V. natriegens DNA |
| | oDS_567_GA_araA_D_fwd | AAAGTATTGGAGTAAGTGGCAAAGAACAGC | oDS_568_GA_araA_D_rev | CTGGTGGAGATGTCAGTGTTTAC | V. natriegens DNA |
| | oDS_570_GA_araA_V_fwd | CGGTAAACACTGACATCTCCACCAGTAGTAGCGGCCGCTGCAGTC | oDS_569_GA_araA_V_rev | TCGCTGCCCTATCTGGCGTAACTATGGCCGCGAATTCCAGAAATC | pMC_V_01 |
| | oDS_565_GA_araA_U_fwd | ATAGTTACGCCAGATAGGGCAG | oDS_688_GA_araA C:C_U_rev_new | TCTTTGCGACTTACTCCAATACTTTAGGACCGTATAGG | V. natriegens DNA |

| | | | | | |
|---|---|---|---|---|---|
| araA C123>T, G124>C tDNA template | oDS_643_GA_araA C:C_U_fwd | AAAGTATTGGAGTAAGTCGCAAAGAACAGCGAAGAG | oDS_568_GA_araA_D_rev | CTGGTGGAGATGTCAGTGTTTAC | V. natriegens DNA |
| | oDS_570_GA_araA_V_fwd | CGGTAAACACTGACATCTCCACCAGTAGTAGCGGCCGCTGCAGTC | oDS_569_GA_araA_V_rev | TCGCTGCCCTATCTGGCGTAACTATGGCCGCGAATTCCAGAAATC | pMC_V_01 |

*Supplementary Table S5: Oligonucleotides to generate tDNA fragments from tDNA template plasmids. All sequences written as 5' -> 3'.*

| tDNA | forward oligonucleotide | | reverse oligonucleotide | | PCR template |
|---|---|---|---|---|---|
| wbfF 3kb | oDS_193_GA_dwbf_U_fwd | TAGCAACTGTTTTAGCGCTGAGC | oDS_196_GA_dwbf_D_rev | GTGTTCCTGTCGATAAGTATTGATC | wbfF tDNA template |
| wbfF 1kb | oDS_410_tDNA_wbf_1kb_U_fwd | GAACGATGTCGACAGCGATG | oDS_411_tDNA_wbf_1kb_D_rev | TCTAAATTGTTGAAATTTTTTTTCAATCATGATG | wbfF tDNA template |
| wbfF 500bp | oDS_412_tDNA_wbf_500bp_U_fwd | ACGAGCTTCATTGAGAAATATATACAC | oDS_413_tDNA_wbf_500bp_D_rev | ATGCAATGACCGCCAGTAATAAG | wbfF tDNA template |
| wbfF 200bp | oDS_414_tDNA_wbf_200bp_U_fwd | TTATCTAGGCGAGTTTTATACGTCAG | oDS_415_tDNA_wbf_200bp_D_rev | TGTCATCGAAAGAACTCTTAATTTCG | wbfF tDNA template |
| wbfF 100bp | oDS_416_tDNA_wbf_100bp_U_fwd | AAAAATCAGCAGAACAACGACAC | oDS_417_tDNA_wbf_100bp_D_rev | GACCTAAAGAACGGGTGGATG | wbfF tDNA template |
| wbfF 50bp | oDS_418_PE_wbf_50bp_U_fwd | TGAAAATGCCTCTTTTTTAAAACTTTAATGAATGAGTATGATGATAAAAGCAATTGATG | oDS_419_PE_wbf_50bp_U_rev | TTGTTTTTGTTAACCCAATTTACGAATAAATGACGCTTATACATCAATTGCTTTTATC | Primer extension |
| wbfF 3kb/50bp | oDS_193_GA_dwbf_U_fwd | TAGCAACTGTTTTAGCGCTGAGC | oDS_616_tDNA_wbf_50bp_rev | TTGTTTTTGTTAACCCAATTTACGAATAAATG | wbfF tDNA template |
| xds | oDS_531_GA_xds_U_fwd | TCGGTTACGTTTTTCAGGAGC | oDS_534_GA_xds_D_rev | TGCCGATTTTCGTCCCGATC | xds tDNA template |
| flagella | oDS_355_GA_flagella_U_fwd | ACTTACAGAATGAGCGTAAATGG | oDS_358_GA_flagella_D_rev | TGTCAACGGCACTTTGATGG | flagella tDNA Template |
| pili | oDS_497_GA_pili_U_fwd | CAGTTGCTCGAAAATCTCAACC | oDS_500_GA_pili_D_rev | TGGTTGGATCTATAAAACAAGCAAATC | pili tDNA template |
| galE | oDS_507_GA_galE_U_fwd | GCAAGGCGCACGATTAAACC | oDS_510_GA_galE_D_rev | ATTTTTTATTGGAACTTGACTGGTTAG | galE tDNA template |
| vnp1 | oDS_541_GA_vnp1_U_fwd | TCGGCACACAAATTGGTACG | oDS_544_GA_vnp1_D_rev | ACCACAACACTCAATTTGGACG | vnp1 tDNA template |
| vnp2 | oDS_553_GA_vnp2_U_fwd | CAAATCCTTCGGACAGCAGAG | oDS_556_GA_vnp2_D_rev | GCAGAGCAACACGCAGCTC | vnp2 tDNA template |
| rpoS-mScarlet-I (indirect) | oDS_366_GA_rpoS_U_fwd | AAGCGCCATTTTGAGTCTGC | oDS_371_GA_rpoS_D_rev | ACTGATCGAGACGCCTATCG | rpoS mScarlet-I tDNA template (indirect) |
| rpoS-mScarlet-I (direct) | oDS_366_GA_rpoS_U_fwd | AAGCGCCATTTTGAGTCTGC | oDS_371_GA_rpoS_D_rev | ACTGATCGAGACGCCTATCG | rpoS mScarlet-I tDNA template (direct) |
| hisG-mScarlet-I-I | oDS_426_GA_hisG_U_fwd | TCACGCCATCTTCACCATCAAAG | oDS_429_GA_hisG_D_rev | CCTTGCTCATCAAGATTCACG | tDNA template hisG-mScarlet-I |
| rplS-mScarlet-I (indirect) | oDS_437_GA_rplS_U_fwd | GATGATGGGTGAGATTAAAGATC | oDS_442_GA_rplS_D_rev | TTATTGTTTTTCTTGGTGTGGTCG | tDNA template rplS-mScarlet-I (indirect) |

| | | | | | |
|---|---|---|---|---|---|
| rplS-mScarlet-I (direct) | oDS_437_GA_rplS_U_fwd | GATGATGGGTGAGATTAAAGATC | oDS_442_GA_rplS_D_rev | TTATTGTTTTTCTTGGTGTGGTCG | tDNA template rplS-mScarlet-I (direct) |
| rpsB-Scarlet-I | oDS_450_GA_rpsB_U_fwd | GATAAAACTCCTTCATCATCATTTCG | oDS_453_GA_rpsB_D_rev | ATACCAGTGTTTTCTGCCGTTAG | tDNA template rpsB-mScarlet-I |
| int9::mScarlet-I | oDS_461_GA_int9_U_fwd | TTTGTTTCGCCAGAGTATGGAG | oDS_464_GA_int9_D_rev | CTCCATGTTTATGTCTAACACGG | int9 mScarlet-I integration tDNA template |
| malQ 298C>T | oDS_391_GA_malQ_U_fwd | CAAATATGGCACTAAGTGTTCAGC | oDS_394_GA_malQ_D_rev | AAAGCTTTGTTGCGATTCAGATC | malQ tDNA Template |
| glpK 76C>T | oDS_587_GA_glpK_U_fwd | CCAAGAATGGCGTTTGCTGC | oDS_590_GA_glpK_D_rev | ATTCAAAACCGGTTTTTATCTCCG | glpK tDNA template |
| glpK 76C>T, G81>C | oDS_587_GA_glpK_U_fwd | CCAAGAATGGCGTTTGCTGC | oDS_590_GA_glpK_D_rev | ATTCAAAACCGGTTTTTATCTCCG | glpK C:C tDNA template |
| araA C123>T | oDS_565_GA_araA_U_fwd | ATAGTTACGCCAGATAGGGCAG | oDS_568_GA_araA_D_rev | CTGGTGGAGATGTCAGTGTTTAC | araA tDNA template |
| araA C123>T, G124>C | oDS_565_GA_araA_U_fwd | ATAGTTACGCCAGATAGGGCAG | oDS_568_GA_araA_D_rev | CTGGTGGAGATGTCAGTGTTTAC | araA C:C tDNA template |

*Supplementary Table S6: Oligonucleotides used for PCR screening of deletions and integrations. All sequences written as 5' -> 3'.*

| Mutation | forward oligonucleotide | | reverse oligonucleotide | |
|---|---|---|---|---|
| Δxds | oDS_537_S_xds_out_fwd | GTTTTTCTTCCTACCTTTGAACG | oDS_538_S_xds_gap_rev | TAAACACTTATGTCCCATGTTTTATAG |
| Δflagella | oDS_363_S_flagella_out_fwd | CTCGAAGCACGCCAAATTGC | oDS_361_S_flagella_gap_rev | ATCATAAAGTCAGTGTTTTAAATATGTGC |
| Δpili | oDS_503_S_pili_out_fwd | ACGCTGTTTTTCAAGATCTTTGG | oDS_504_S_pili_gap_rev | AGAAGGAAGAGAGATTTCACTTTG |
| ΔgalE | oDS_513_S_galE_out_fwd | TTGGCTCCGGTAATCAAACG | oDS_514_S_galE_gap_rev | AGCGGAGAAGTAAAATTTTCATCC |
| Δvnp1 | oDS_547_S_vnp1_out_fwd | TCTGCTAGGTTTGCACAAGC | oDS_548_S_vnp1_gap_rev | ATGTAAAAACCACACAAACAAATCTAC |
| Δvnp2 | oDS_559_S_vnp2_out_fwd | GATGGTTTAGGCTCCAATACC | oDS_560_S_vnp2_gap_rev | GTAGTCAAAGGAAGAAAAAAGTGG |
| rpoS-mScarlet-I (indirect) | oDS_377_S_rpoS_out_fwd | CTCGGTGTTCAGTCTGTTGC | oDS_375_GA_mScarlet_rev | TTTGTACAGTTCATCCATACCAC |
| rpoS-mScarlet-I (direct) | oDS_377_S_rpoS_out_fwd | CTCGGTGTTCAGTCTGTTGC | oDS_375_GA_mScarlet_rev | TTTGTACAGTTCATCCATACCAC |
| hisG-mScarlet-I | oDS_432_S_hisG_out_fwd | CACTTCTTGAGTCATAGTGTACG | oDS_375_GA_mScarlet_rev | TTTGTACAGTTCATCCATACCAC |
| rplS-mScarlet-I (indirect) | oDS_445_S_rplS_out_fwd | CGACGTTCTGATTGTCGATACC | oDS_375_GA_mScarlet_rev | TTTGTACAGTTCATCCATACCAC |
| rplS-mScarlet-I (direct) | oDS_445_S_rplS_out_fwd | CGACGTTCTGATTGTCGATACC | oDS_375_GA_mScarlet_rev | TTTGTACAGTTCATCCATACCAC |
| rpsB-Scarlet-I | oDS_456_S_rpsB_out_fwd | AGCCGCGTCGCTGAAAATCG | oDS_375_GA_mScarlet_rev | TTTGTACAGTTCATCCATACCAC |
| int9::mScarlet-I | oDS_469_S_int9_out_fwd | GCATTGCAAAGCTGGCAACC | oDS_375_GA_mScarlet_rev | TTTGTACAGTTCATCCATACCAC |

*Supplementary Table S7: Oligonucleotides used for PCR to amplify target region for sequencing. Oligonucleotide in right column is nested primer used for Sanger sequencing. All sequences written as 5' -> 3'.*

| Mutation | forward oligonucleotide | | reverse oligonucleotide | | oligonucleotide for sequencing | |
|---|---|---|---|---|---|---|
| Δxds | oDS_537_S_xds_out_fwd | GTTTTTCTTCCTACCTTTGAACG | oDS_539_S_xds_D_rev | AGCCAAGCGATAAAAAGCCTTG | oDS_540_S_xds_U_fwd | CATCTTTGCACCAAATGTGTGC |
| Δflagella | oDS_363_S_flagella_out_fwd | CTCGAAGCACGCCAAATTGC | oDS_364_S_flagella_D_rev | GTTAAGTCTTGGAAGTCTTGTGC | oDS_365_S_flagelle_U_fwd | ACGTTGGCCGTGTAGACATC |
| Δpili | oDS_503_S_pili_out_fwd | ACGCTGTTTTTCAAGATCTTTGG | oDS_505_S_pili_D_rev | CATCTCTTATCCCTGAAGATGC | oDS_506_S_pili_U_fwd | TCACGGTAAGTGGCCAGTCG |
| ΔgalE | oDS_513_S_galE_out_fwd | TTGGCTCCGGTAATCAAACG | oDS_515_S_galE_D_rev | GGAACGTAACTGAAGGTTCG | oDS_516_S_galE_U_fwd | CTTATTCTCAAACGCTTGGACC |
| Δvnp1 | oDS_547_S_vnp1_out_fwd | TCTGCTAGGTTTGCACAAGC | oDS_549_S_vnp1_D_rev | GAGGTATGAATATGAGTGCTGG | oDS_550_S_vnp1_U_fwd | GAAGACTTAGCGCAGATAAAGC |
| Δvnp2 | oDS_559_S_vnp2_out_fwd | GATGGTTTAGGCTCCAATACC | oDS_561_S_vnp2_D_rev | AGGCTGATCAGCTTTTCTCG | oDS_562_S_vnp2_U_fwd | TCCTTATAGTGATATGCGATCAG |
| rpoS-mScarlet-I (indirect) | oDS_377_S_rpoS_out_fwd | CTCGGTGTTCAGTCTGTTGC | oDS_378_S_rpoS_D_rev | AACTCTAAGTCAAAAGCACTCG | oDS_379_S_rpoS_U_fwd | CGATTCACGTTGTGAAAGAGC |
| rpoS-mScarlet-I (direct) | oDS_377_S_rpoS_out_fwd | CTCGGTGTTCAGTCTGTTGC | oDS_378_S_rpoS_D_rev | AACTCTAAGTCAAAAGCACTCG | oDS_379_S_rpoS_U_fwd | CGATTCACGTTGTGAAAGAGC |
| hisG-mScarlet-I | oDS_432_S_hisG_out_fwd | CACTTCTTGAGTCATAGTGTACG | oDS_433_S_hisG_D_rev | CCTGCGTTATCAAGCAGAGG | oDS_434_S_hisG_U_fwd | TGCTACGACTTATCCGCACC |
| rplS-mScarlet-I (indirect) | oDS_445_S_rplS_out_fwd | CGACGTTCTGATTGTCGATACC | oDS_446_S_rplS_D_rev | TCAGGCACTGAAAGATGATCG | oDS_447_S_rplS_U_fwd | CGAACTTGGCTAAGAAGACC |
| rplS-mScarlet-I (direct) | oDS_445_S_rplS_out_fwd | CGACGTTCTGATTGTCGATACC | oDS_446_S_rplS_D_rev | TCAGGCACTGAAAGATGATCG | oDS_447_S_rplS_U_fwd | CGAACTTGGCTAAGAAGACC |
| rpsB-Scarlet-I | oDS_456_S_rpsB_out_fwd | AGCCGCGTCGCTGAAAATCG | oDS_457_S_rpsB_D_rev | GCTCATAAAAAGACCGCAGC | oDS_458_S_rpsB_U_fwd | TACTAAGCGCGCTGCATCTG |
| int9::mScarlet-I | oDS_469_S_int9_out_fwd | GCATTGCAAAGCTGGCAACC | oDS_472_S_int9_D_rev | AGGAAAAACTGGCGTTGAGC | oDS_471_S_int9_U_fwd | CTTACGCTGGTTACAACAGC |
| malQ 298C>T | oDS_403_S_malQ_out_fwd | GTTCAACGTGGGTGTACTCC | oDS_404_S_malQ_D_rev | GAGACTATGTGAACAACATCTGG | oDS_405_S_malQ_U_fwd | CGAATTCTAAGAATGCACGTGC |
| glpK 76C>T, G81>C | oDS_593_S_glpK_out_fwd | CTCCATCCCAACCAAAGTGG | oDS_594_S_glpK_D_rev | CACGAATAATATGGTTTGAGTTAACG | oDS_595_S_glpK_U_fwd | CAATGAACCCACTATTGATTGG |
| araA C123>T, G124>C | oDS_571_S_araA_out_fwd | CATTTGAACCGAAAAATGCGTTGG | oDS_572_S_araA_D_rev | CTCAATTGATGGCTTGTCAGC | oDS_573_S_araA_U_fwd | GCAGCGATTCCAGCCTTTGG |
